# mRNA stability fine tunes gene expression in the developing cortex to control neurogenesis

**DOI:** 10.1101/2024.07.22.604643

**Authors:** Lucas D. Serdar, Jacob R. Egol, Brad Lackford, Brian D. Bennett, Guang Hu, Debra L. Silver

**Affiliations:** Department of Molecular Genetics and Microbiology, Duke University Medical Center, Durham, NC 27710; National Institute of Environmental Health Sciences, Durham, NC 27709; Departments of Cell Biology and Neurobiology, Duke University Medical Center, Durham, NC 27710; Duke Institute for Brain Sciences and Duke Regeneration Center, Duke University Medical Center, Durham, NC 27710

**Keywords:** RNA stability, cortical development, CNOT3, CCR4-NOT, slam seq, neurogenesis

## Abstract

RNA expression levels are controlled by the complementary processes of synthesis and degradation. Although mis-regulation of RNA turnover is linked to neurodevelopmental disorders, how it contributes to cortical development is largely unknown. Here, we profile the RNA stability landscape of the cortex across development and demonstrate that control of stability by the CCR4-NOT complex is essential for corticogenesis *in vivo*. First, we use SLAM-seq to measure RNA half-lives transcriptome-wide across multiple stages of cortical development. We characterize *cis*-acting features associated with RNA stability and find that RNAs that are upregulated across development tend to be more stable, while downregulated RNAs are less stable. To probe how disruption of RNA turnover impacts cortical development, we assess developmental requirements of CNOT3, a core component of the CCR4-NOT deadenylase complex. Mutations in *CNOT3* are associated with human neurodevelopmental disorders, however its role in cortical development is unknown. Conditional knockout of *Cnot3* in neural progenitors and their progeny in the developing mouse cortex leads to severe microcephaly due to reduced neuron production and p53-dependent apoptosis. Collectively, our findings demonstrate that fine-tuned control of RNA turnover is crucial for brain development.

## Introduction

The cerebral cortex is essential for higher order functions, including cognitive reasoning, and somatosensory, motor, and visual processing. These processes rely on proper embryonic development, and defective embryonic corticogenesis can lead to neurodevelopmental disorders, including autism, schizophrenia, and intellectual disability. The developmental trajectory and underlying biological and molecular events necessary to construct the cortex during development are generally well defined (Greig et al. 2013; Villalba, Gotz and Borrell 2021). In mice, the main neural progenitors are radial glial cells (RGCs), which produce neurons directly, and intermediate progenitors (IPs), which are derived from RGCs and are also neurogenic (Fig. 1A). The mature mammalian cortex consists of a stereotypical six-layered architecture that forms in an inside-out fashion. Neurons born early in development (∼embryonic day (E)12.5 – E13.5 in mouse) occupy deep layers, while late born neurons (∼E14.5 – E16.5) typically occupy superficial layers (Di Bella et al. 2024). Thus, corticogenesis is a dynamic process in which progenitor potency and neuronal composition changes with development.

**Figure 1.**
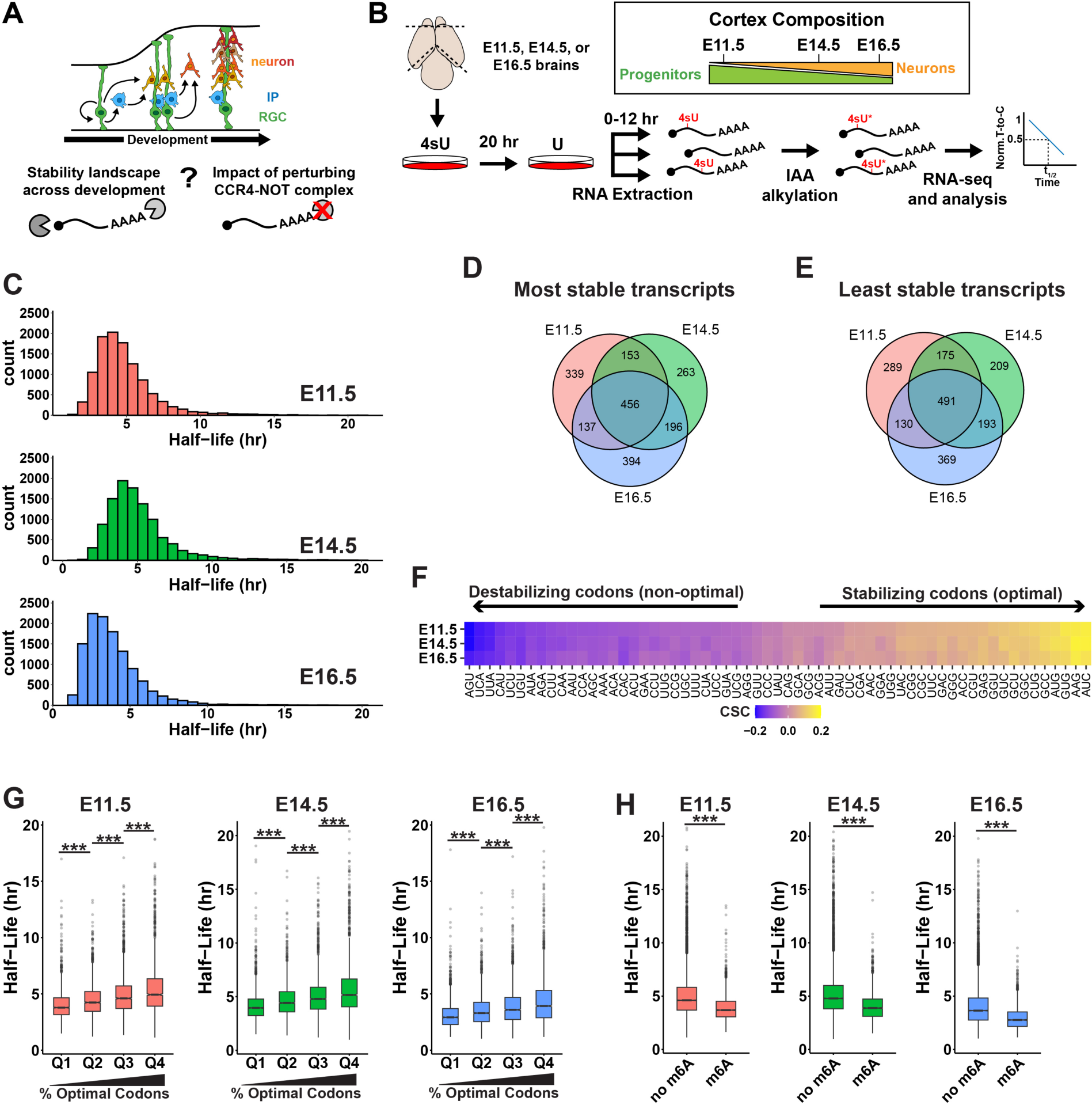
Features of mRNA stability across cortical development. (A) Overview of cortical development showing radial glial cells (RGC) generating neurons directly, or indirectly through intermediate progenitors (IP), with goals of this study schematized below. (B) Approach for SLAM-seq experiment, with relative cellular composition of the cortex at the three developmental stages used in this study (n = 3 biological replicates per stage). (C) Histograms showing distribution of RNA half-lives. (D and E) Venn diagrams for top 10% most and least stable transcripts identified at each stage. (F) Codon stabilization coefficient (CSC) values for each of the 61 non-stop codons. (G) Half-lives of RNAs binned into quartiles based on percentage of optimal (positive CSC) codons. (H) Half-lives of RNAs with or without m^6^A. *** *p* < 0.001. One-way ANOVA with Tukey’s HSD post-hoc test (J), Wilcoxon rank-sum test (K).

In line with diverse cellular processes, the transcriptomes of progenitor and neurons are dynamic during corticogenesis (Fietz et al. 2012; Molyneaux et al. 2015; Telley et al. 2019; Di Bella et al. 2021). These developmental gene expression changes are controlled by transcriptional and post-transcriptional mechanisms (Kwan, Sestan and Anton 2012; Lennox, Mao and Silver 2018). RNA stability regulation is emerging as an important modulator of gene expression during cortical development. Previous work has revealed roles for core components of the RNA decay machinery, including the exosome (Ulmke et al. 2021) and the nonsense-mediated RNA decay (NMD) pathway (Tarpey et al. 2007; Lou et al. 2014; Huang et al. 2018; Jaffrey and Wilkinson 2018; Kurosaki et al. 2021; Lin et al. 2024). Several studies have also highlighted the importance of additional *cis*- and *trans*-acting regulators of stability in cortical development. For example, the oscillatory RNA expression pattern of the Notch signaling pathway gene *Hes1* in cortical progenitors is maintained in part by microRNA-mediated degradation (Hirata et al. 2002; Bonev, Stanley and Papalopulu 2012). In addition, the abundant epitranscriptomic mark N6-methyladenosine (m^6^A) destabilizes RNA in cortical cells and influences neurogenesis (Yoon et al. 2017). While these studies highlight fundamental roles of mRNA stability in cortical development, our understanding of this process is limited. Indeed, a survey of RNA half-lives across development is lacking, as is a mechanistic understanding of how RNA turnover impacts the developing brain.

A key regulator of mRNA stability is the CCR4-NOT deadenylase complex (Collart 2016). This multi-subunit complex catalyzes the removal of 3’ poly(A) tails from mRNAs, typically one of the first steps in mRNA degradation. CNOT3 is a non-enzymatic component of the CCR4-NOT complex, but is required for efficient deadenylation and mRNA degradation (Boland et al. 2013; Stowell et al. 2016; Raisch et al. 2019). Mutations in *CNOT3* as well as another component, *CNOT1*, are linked to neurodevelopmental pathologies characterized by a spectrum of intellectual disability, speech delay, seizures, and behavioral problems (Kruszka et al. 2019; Martin et al. 2019; Meyer et al. 2020; Vissers et al. 2020; Priolo et al. 2021; Lv et al. 2023; Zhao et al. 2023). Of note, *CNOT3* is categorized as a “High-confidence, syndromic” autism risk gene in the SFARI autism gene database. Previous work implicates CNOT3 in diverse stem cell populations, including cardiomyocytes (Zhou et al. 2017; Yamaguchi et al. 2018), embryonic stem cells (Zheng et al. 2012; Zheng et al. 2016), and spermatogonial stem cells (Chen et al. 2023). Despite clinical significance of *CNOT3* in neurodevelopment, its molecular function in cortical development is entirely unknown.

Here, we employ multi-pronged approaches to understand the principles of RNA degradation in the developing brain and how its misregulation impacts corticogenesis (Fig. 1A). First, we use SLAM-seq to globally measure RNA half-lives across multiple stages of mouse development. From this we identify key features governing RNA stability in the cortex and uncover relationships between RNA turnover rates and gene expression changes across development. Second, we employ *Cnot3* conditional knockout mouse models to demonstrate its essential role in cortical neurogenesis. Our findings reveal the importance of RNA stability control in the developing cortex and establish the CCR4-NOT complex as a key factor contributing to proper gene expression and cellular function during corticogenesis.

## Results

### SLAM-seq defines the landscape and features of RNA stability in cortical cells

To characterize the mRNA stability landscape of the developing cortex, we performed thiol(SH)-linked alkylation for metabolic sequencing (SLAM-seq) (Herzog et al. 2017) at three different developmental stages. We chose E11.5, E14.5, and E16.5 as representative stages of early, middle, and late neurogenesis, respectively. E11.5 cortices are predominantly composed of RGCs, while E14.5 and E16.5 cortices contain increasing numbers of intermediate progenitors and neurons, along with non-RGC derived cells. Cortices from E11.5, E14.5, and E16.5 embryos were dissociated into single cell suspensions using three independent biological replicates per stage and cultured *in vitro* using conditions permissive for proliferation (Shen et al. 2002). 4-thiouridine (4sU) was added to culture media for 20 hr to label nascent RNAs. Cells were then collected for RNA extraction 0, 2, 4, 8, and 12 hr following a chase with excess unmodified uridine to monitor degradation of labeled RNAs over time (Fig. 1B). Following sequencing, we obtained reproducible half-life measurements for 10,842 transcripts at E11.5, 10,671 transcripts at E14.5, and 11,832 transcripts at E16.5 (Fig. 1C, Supplemental Table S1). Principal component analysis (PCA) showed that the biological replicates within each stage largely clustered together but were distinct between the stages (Supplemental Fig. S1A). Comparison of z-score normalized half-lives revealed high correlations between stages (r > 0.87), indicating that the relative stability of individual transcripts is highly consistent across cortical development (Supplemental Fig. S1B, C).

To further compare the RNA stability landscape between stages, we assessed the top 10% most and least stable transcripts at E11.5, E14.5, and E16.5. We observed a high degree of similarity between the three stages. Among the most stable transcripts, 456 were shared among all three stages. Similarly, 491 of the least stable transcripts were shared across all three stages (Fig. 1E). Notably, more transcripts were shared between all three stages than were unique to each stage. Gene ontology (GO) analysis of the most and least stable transcripts at each stage revealed consistently enriched terms shared across developmental stages (Supplemental Fig. S1D, E). Stable transcripts were enriched for functions such as ATP synthesis and protein synthesis, while unstable genes encoded regulators of transcription and cell fate determination. Importantly, these functional classes for stable and unstable transcripts are similar to those observed in other cell types in diverse organisms (Thomsen et al. 2010; Miller et al. 2011; Schwanhausser et al. 2011; Burow et al. 2015; Herzog et al. 2017).

We next sought to identify *cis*-acting factors that shape RNA turnover dynamics in cortical cells. We first focused on 5′ and 3′ UTRs as they are known to contain regulatory elements important for RNA stability (Mayr 2019; Jia et al. 2020). We examined 3′ UTR sequences of the same top 10% most stable and least stable transcripts at each stage (Engel et al. 2022). The most stable RNAs had shorter, more GC-rich, 3′ UTRs (Supplemental Fig. S1F, G). Analysis of 5′ UTR sequences revealed a similar relationship between length and stability, with stable transcripts having shorter 5′ UTRs (Supplemental Fig. S1H). However, no relationship was observed between 5′ UTR GC content and stability (Supplemental Fig. S1I). The preferential impact of 3′ UTR GC content on RNA stability may reflect the enrichment of regulatory elements within these regions (Caput et al. 1986).

Codon content has also been implicated in mRNA stability regulation in multiple biological contexts (Presnyak et al. 2015; Bazzini et al. 2016; Burow et al. 2018; Narula et al. 2019; Wu et al. 2019; Forrest et al. 2020). To determine whether mRNA stability in the developing mouse cortex is associated with codon content, we calculated the codon stabilization coefficient (CSC), a measure of the correlation between codon usage and mRNA half-life (Presnyak et al. 2015). We found robust correlations between codon usage and mRNA stability across all three stages, with CSC values ranging from 0.17 (stabilizing/optimal) to -0.2 (destabilizing/non-optimal) (Fig. 1F). CSC values for each codon were virtually identical between E11.5, E14.5, and E16.5 (Supplemental Fig. S1J, K). To further assess the impact of codon content on mRNA stability, we binned mRNAs into quartiles based on their optimal codon content and then compared the half-lives of each quartile. We found a clear and consistent trend across all three developmental stages showing that mRNAs with a higher proportion of optimal codons are more stable (Fig. 1G).

Finally, we examined the impact of m^6^A, an abundant epitranscriptomic mark associated with multiple aspects of mRNA metabolism, including mRNA stability (Lee et al. 2020). m^6^A was previously shown to be present in >2,000 transcripts in E14.5 mouse primary cortical cells, where it was implicated in RNA degradation (Yoon et al. 2017). To assess the destabilizing impact of m^6^A in our dataset, and to determine whether its role changes across development, we compared half-lives of mRNAs predicted to contain m^6^A to those without m^6^A at each stage (Yoon et al. 2017) (Fig. 1H). This analysis showed significantly lower half-lives for m^6^A-containing transcripts, with similar extents of destabilization at all stages. This strongly implicates m^6^A in RNA stability throughout neurogenesis, reinforcing previous findings (Yoon et al. 2017). Collectively, our data highlight multiple *cis*-acting features that shape RNA stability in cortical cells.

Although we did not observe widespread shifts in RNA half-lives across development, some variation among individual transcripts was observed (Supplemental Fig. S1B, C). We therefore aimed to identify transcripts with developmentally regulated RNA half-lives. For this, we employed a regression-based method (Nueda, Tarazona and Conesa 2014) to assess RNA degradation kinetics across time in our SLAM-seq data. We first filtered the data to include only the 9,490 transcripts expressed at all three stages, then converted half-lives to z-scores to normalize the data between stages and allow for direct comparisons. Regression analysis was then used to identify transcripts with significant changes in stability across development. Consistent with our previous observations, half-lives for most transcripts were unchanged. However, a small subset of 294 transcripts (3.1%) had significant differences in stability across all three stages assayed. Hierarchical clustering of this subset based on their half-life dynamics identified 7 distinct clusters (Fig. 2A, and Supplemental Table S2). The temporal half-life profile for each cluster was unique (Fig. 2B). Stability of cluster 2 and 3 RNAs gradually increased, while the stability of cluster 5 and 7 RNAs gradually decreased. Cluster 1 and cluster 4 stability was greatest at E16.5 and E14.5, respectively, while cluster 6 transcripts were least stable at E14.5. Altogether, these data demonstrate that turnover rates for individual transcripts are on average constant across cortical development but are dynamically regulated for a subset of transcripts.

**Figure 2.**
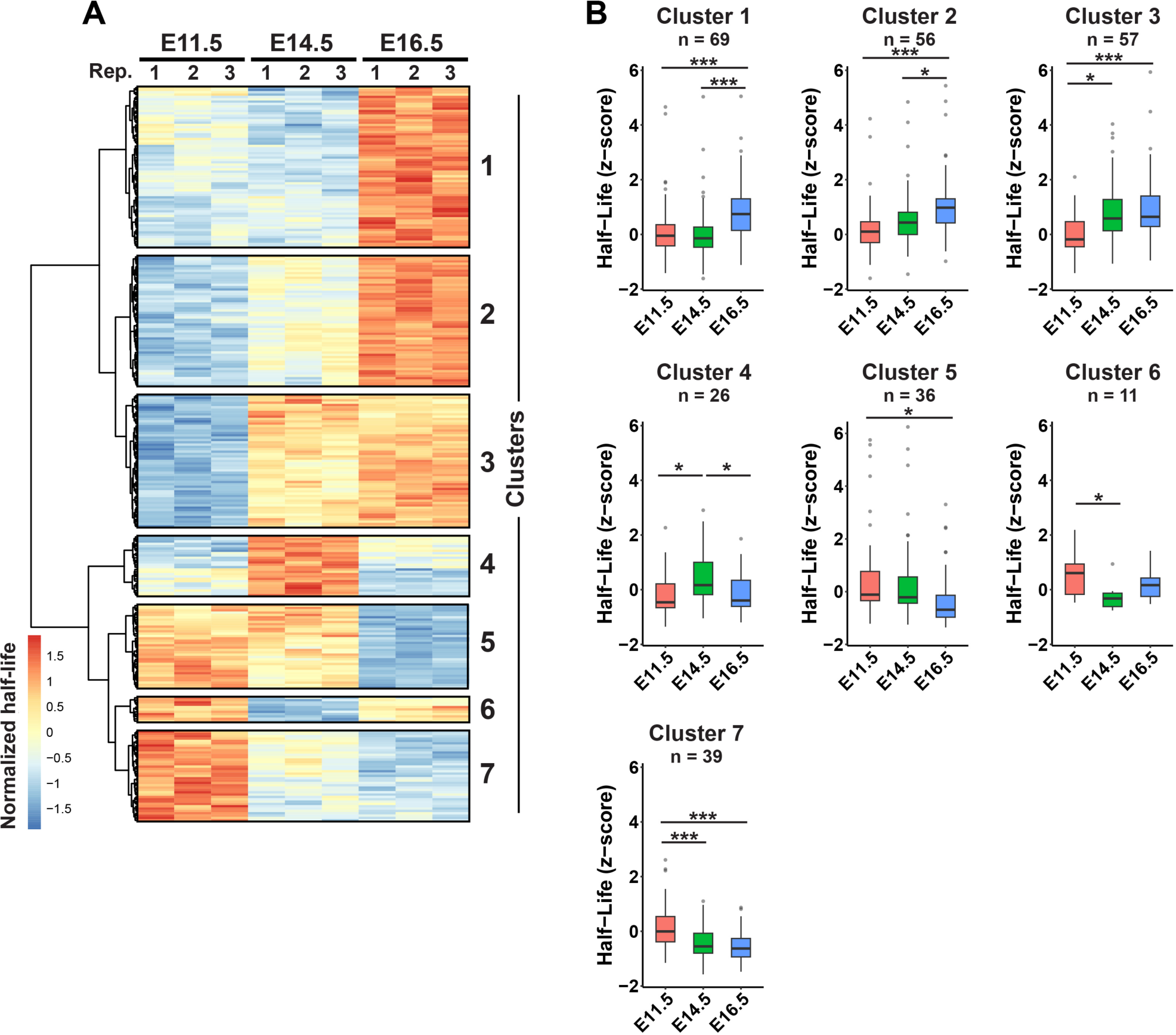
Dynamic patterns of mRNA half-lives across cortical development. (A) Heatmap showing z-score normalized half-lives for 294 significantly changing transcripts at E11.5, E14.5, and E16.5 (n = 3 biological replicates per stage). Numbers to the right indicate cluster labels. (B) Z-score normalized half-lives for transcripts found in the indicated clusters. \**p* < 0.05, \*\**p* < 0.01, \*\*\**p* < 0.001. One-way ANOVA with Tukey’s HSD post-hoc test.

### RNA stability correlates with developmental gene expression changes

Neural progenitors dynamically remodel their transcriptomes over developmental time and upon differentiation (Telley et al. 2019). Yet the extent to which RNA stability regulation influences these transcriptional dynamics is largely unknown. To investigate this question, we assessed the stability of RNAs whose expression significantly changes across development. Because SLAM-seq is conducted using standard RNA-seq, our dataset measures RNA expression levels at each stage, in addition to their half-lives. Using this data, we evaluated differential expression across the three stages. Using cutoffs of fold change > 2 and *p_adj_* < 0.05, we identified 1,056 upregulated and 997 downregulated transcripts at E14.5 compared to E11.5 (Fig. 3A, and Supplemental Table S3). Transcripts that were upregulated from E11.5 to E14.5 included neuronal markers (e.g. *Neurod2, Satb2, Dcx*), while downregulated transcripts included progenitor markers (e.g. *Sox2, Nes, Hes5*). These gene expression changes may reflect the increased abundance of neurons at later stages of development. We further identified 962 upregulated and 730 downregulated genes at E16.5 compared to E14.5 (Fig. 3B, and Supplemental Table S3). Similar to changes from E11.5 to E14.5, we observed increased expression at E16.5 of neuronal markers (e.g. *Satb2, Cux2, Pou3f3/Brn1*) and decreased expression of progenitor and cell cycle markers (*Ccnd2, Sox2, Btg2*). Notably, these data are consistent with transcriptional dynamics that are expected to occur *in vivo* during cortical development.

**Figure 3.**
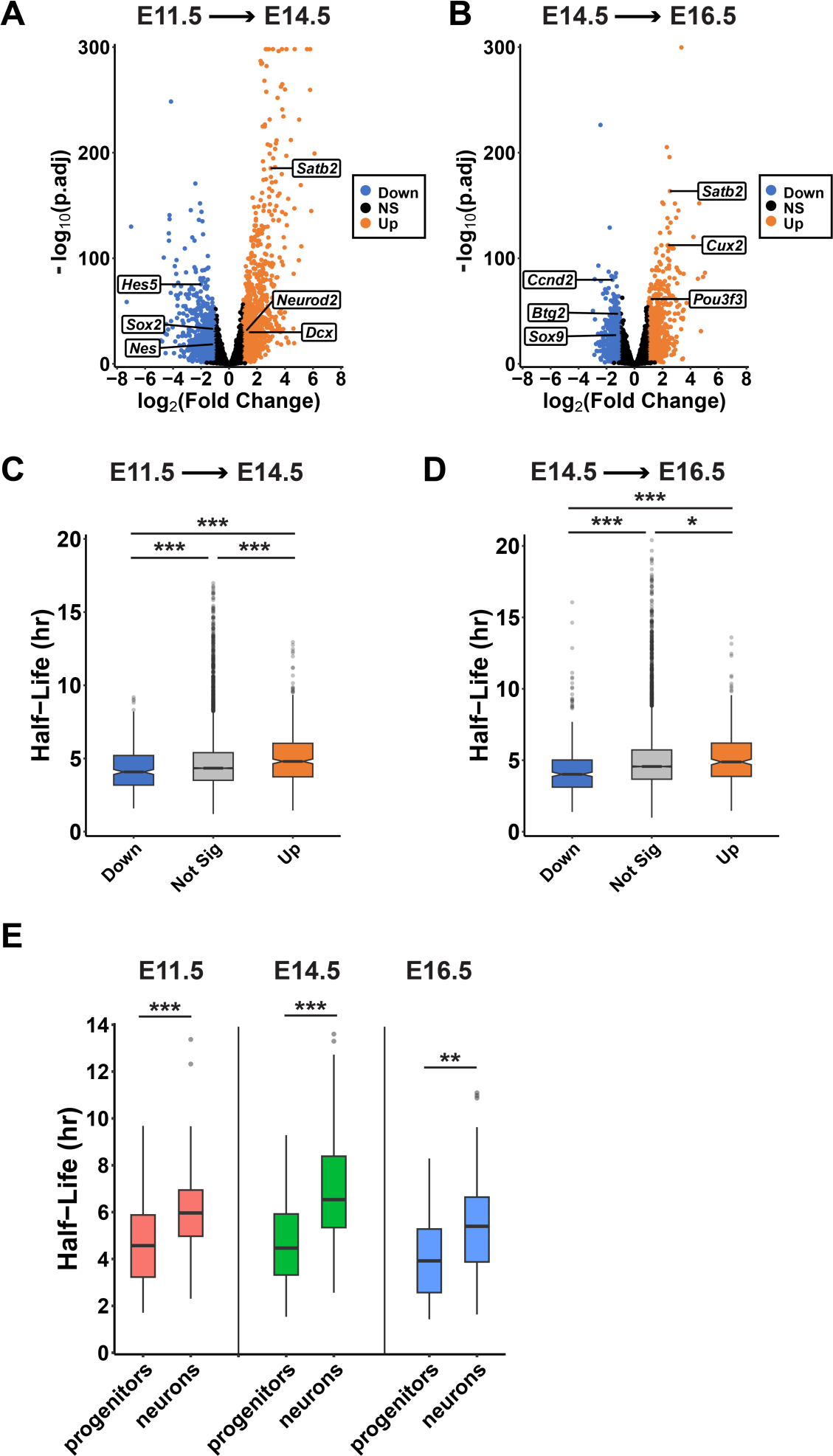
RNA stability correlates with developmental gene expression changes. (A, B) Volcano plots showing transcript expression changes at E14.5 compared to E11.5 (A), or E16.5 compared to E14.5 (B). n = 3 biological replicates per stage. (C, D) Half-lives of differentially expressed transcripts at the indicated developmental transitions. (E) Half-lives for a subset of transcripts enriched in either progenitors (n = 56) or neurons (n = 57). \*\**p* < 0.01, \*\*\**p* < 0.001. One-way ANOVA with Tukey’s HSD post-hoc test (C, D), Wilcoxon rank-sum test (E).

To determine whether there is a relationship between RNA stability and gene expression dynamics across development, we compared half-lives of differentially expressed transcripts. We found that RNAs whose expression is significantly upregulated at E14.5 compared to E11.5 were marginally, but significantly, more stable compared to those that were downregulated (Fig. 3C). The same trend was observed for RNAs that were differentially expressed at E16.5 compared to E14.5 (Fig. 3D). These data suggest that expression dynamics and RNA stability are coordinated. Developmentally downregulated transcripts undergo rapid turnover, while upregulated mRNAs are degraded more slowly to facilitate their accumulation.

Some of these developmentally regulated transcripts are known to have cell-specific expression. Given this, we investigated whether this held true across more of the transcriptome. For this, we mined a previously published scRNA-seq dataset of the developing cortex (Di Bella et al. 2021) to assess expression levels of developmentally regulated transcripts in specific cell types. At both transitions (E11.5 to E14.5 and E14.5 to E16.5), transcripts with decreasing expression were enriched in progenitors (RGCs and IPs), while those with increasing developmental expression were enriched in neurons (Supplemental Fig. S2A, B). Together this suggests that gene expression differences in our datasets are linked to changes in cell composition across development.

Our data demonstrate that some developmentally regulated transcripts exhibit biased cell-type expression profiles (Supplemental Fig. S2A, B) and distinct stability profiles (Fig. 3C, D). Therefore, we next sought to determine whether transcripts enriched in progenitors and neurons have distinct RNA half-lives. To address this, we generated a stringent list of transcripts that are highly enriched in either progenitors (n = 56), or neurons (n = 57) across two independent scRNA-seq datasets (Telley et al. 2019; Di Bella et al. 2021) (Supplemental Table S4). In our SLAM-seq dataset, these transcripts were expressed at all three stages, with progenitor-enriched transcripts most highly expressed at E11.5, and neuron-enriched transcripts most highly expressed at E16.5 (Supplemental Fig. S2C, D). These expression patterns are consistent with known cell composition at each stage. Notably, progenitor-enriched transcripts were significantly less stable than neuron-enriched transcripts (Fig. 3E). This was true across all three developmental stages. Collectively, these data demonstrate cell-specific layers of RNA stability during cortical development and strongly suggest that RNA stability influences developmental expression changes for a subset of transcripts. These data indicate developmental roles of RNA stability for shaping the transcriptome.

### Conditional knockout of *Cnot3* leads to microcephaly and disrupted neuronal lamination

Our data thus far indicate that RNA stability regulation is a key feature of cortical development, implicated in gene expression changes across stages. This led us to investigate the *in vivo* consequences of mis-regulating RNA stability during cortical development. To address this, we focused on CNOT3, a central component of the CCR4-NOT complex (Fig. 4A). CNOT3 was a strong candidate as it is essential for deadenylation (Boland et al. 2013; Stowell et al. 2016; Raisch et al. 2019) and is linked to neurodevelopmental pathologies (Martin et al. 2019; Meyer et al. 2020; Priolo et al. 2021; Lv et al. 2023; Zhao et al. 2023). Importantly, inspection of scRNA-seq datasets of the developing mouse cortex revealed broad *Cnot3* expression in both progenitors and neurons (Telley et al. 2019).

**Figure 4.**
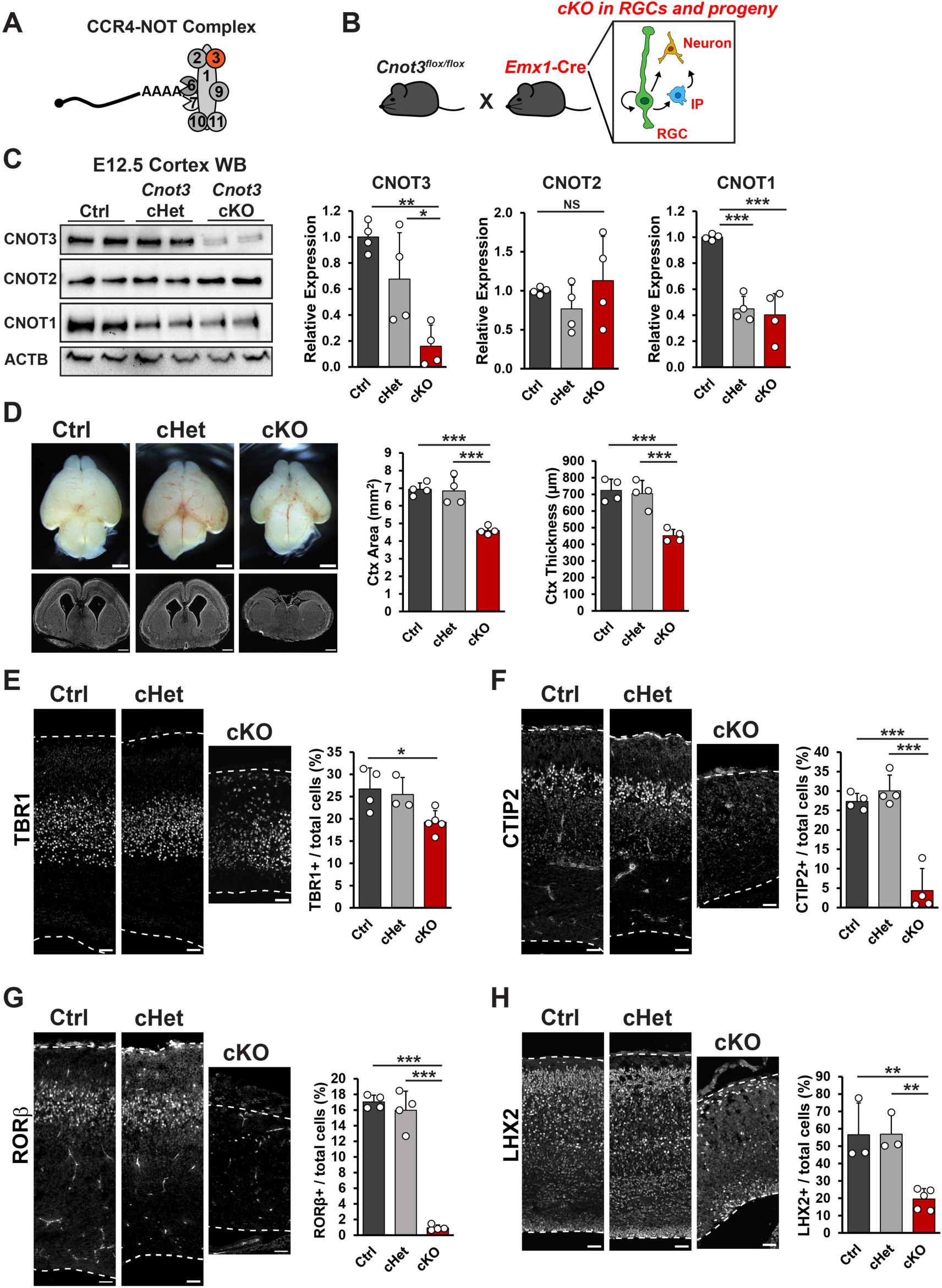
Conditional knockout of *Cnot3* in cortical progenitors impairs brain size and neuronal number. (A) Schematic of CCR4-NOT complex with CNOT3 indicated in red. (B) Conditional knockout (cKO) strategy using *Emx1*-Cre to target radial glial cells (RGC) and their progeny, including intermediate progenitors (IP) and neurons. (C) Western blot using whole cortex protein lysates from E12.5 control (ctrl), conditional heterozygous (cHet), or cKO embryos. Quantifications shown to the right (n = 4 embryos per genotype). (D) Whole mount images and coronal cross sections of E18.5 brains. Quantifications of cortex area and thickness shown to the right (n = 4 embryos per genotype). (E-H) Immunofluorescence for indicated neuronal markers in E18.5 cortical sections, with quantifications shown to the right. n = 3-5 embryos per genotype. \**p* < 0.05, \*\**p* < 0.01, \*\*\**p* < 0.001. One-way ANOVA with Tukey’s HSD post-hoc test. Error bars represent standard deviation. Scale bars: 1 mm (D, top), 500 µm (D, bottom), 50 µm (E-H).

We generated conditional knockouts (cKO) of *Cnot3* in RGCs and their progeny by crossing *Cnot3^lox/lox^* mice with *Emx1*-Cre (Gorski et al. 2002) (Fig. 4B). At E12.5 (∼3 days after the onset of Cre-mediated recombination), we observed an 84% and 82% reduction in CNOT3 protein and RNA expression, respectively, in cKO cortices (Fig. 4C, and Supplemental Fig. S3A). Interestingly, expression of CNOT1 was also reduced. Reduction of CNOT1 protein likely occurs via translational and/or post-translational mechanisms, as we observed no significant change in *Cnot1* RNA measured by RT-qPCR (Supplemental Fig. S3A). No statistically significant changes were observed for CNOT2 expression at either the protein or RNA levels. As CNOT1 is essential for scaffolding the complex, this suggests that the integrity of the entire CCR4-NOT complex may be impaired in this mouse model. These data demonstrate the *Cnot3* cKO can be faithfully used to assess roles of this complex in cortical development.

To determine the requirement of *Cnot3* for cortical development, we first examined brains at E18.5, when neurogenesis is largely complete. cKO mice exhibited profound microcephaly, with a 38% reduction in cortical area, and a similar 38% reduction in cortical thickness (Fig. 4D). In contrast, conditional heterozygous (cHet) brain sizes were comparable to control. This indicates that CNOT3 is required for overall cortical size. The cortex forms a six-layered laminar architecture, with early born neurons present in deep layers and later born neurons primarily present in superficial layers (Di Bella et al. 2024). To examine the extent to which *Cnot3* ablation affects glutamatergic projection neurons, we performed immunostaining for TBR1 (Layer VI), CTIP2 (Layer V), RORβ (Layer IV), and LHX2 (Layer II/III). In cKO brains compared to control, we observed a ∼28% reduction in TBR1+ neuron density, a near complete loss of CTIP2+ and RORβ+ neurons, and a 65% reduction in LHX2+ neurons (Fig. 4E-H). cHet brains were similar to control. In addition to reduced density, neuronal distribution for some markers was also disrupted in cKO brains (Supplemental Fig. S3B-E). These data indicate that CNOT3 is essential for number and laminar position of glutamatergic neurons produced throughout cortical development.

### *Cnot3* is required for viability of both progenitors and neurons

To understand how *Cnot3* controls brain size and neuron number, we next examined its requirements for neurogenesis. At E14.5, mid-neurogenesis, we observed an 18% reduction in cortical thickness in *Cnot3* cKO mice, indicating a role for *Cnot3* in development at earlier stages (Fig. 5A, B). We performed immunofluorescence against SOX2 and PAX6 to assess RGCs. Both markers were qualitatively reduced in the proliferative zones (Supplemental Fig. S4A). Indeed, we observed a 26% reduction in apical SOX2+ RGCs. We also assessed IPs using TBR2 expression. This showed a 43% reduction in IPs in cKO cortices compared to control (Fig. 5C, D). This demonstrates that *Cnot3* is critical for proper number of both RGCs and IPs.

**Figure 5.**
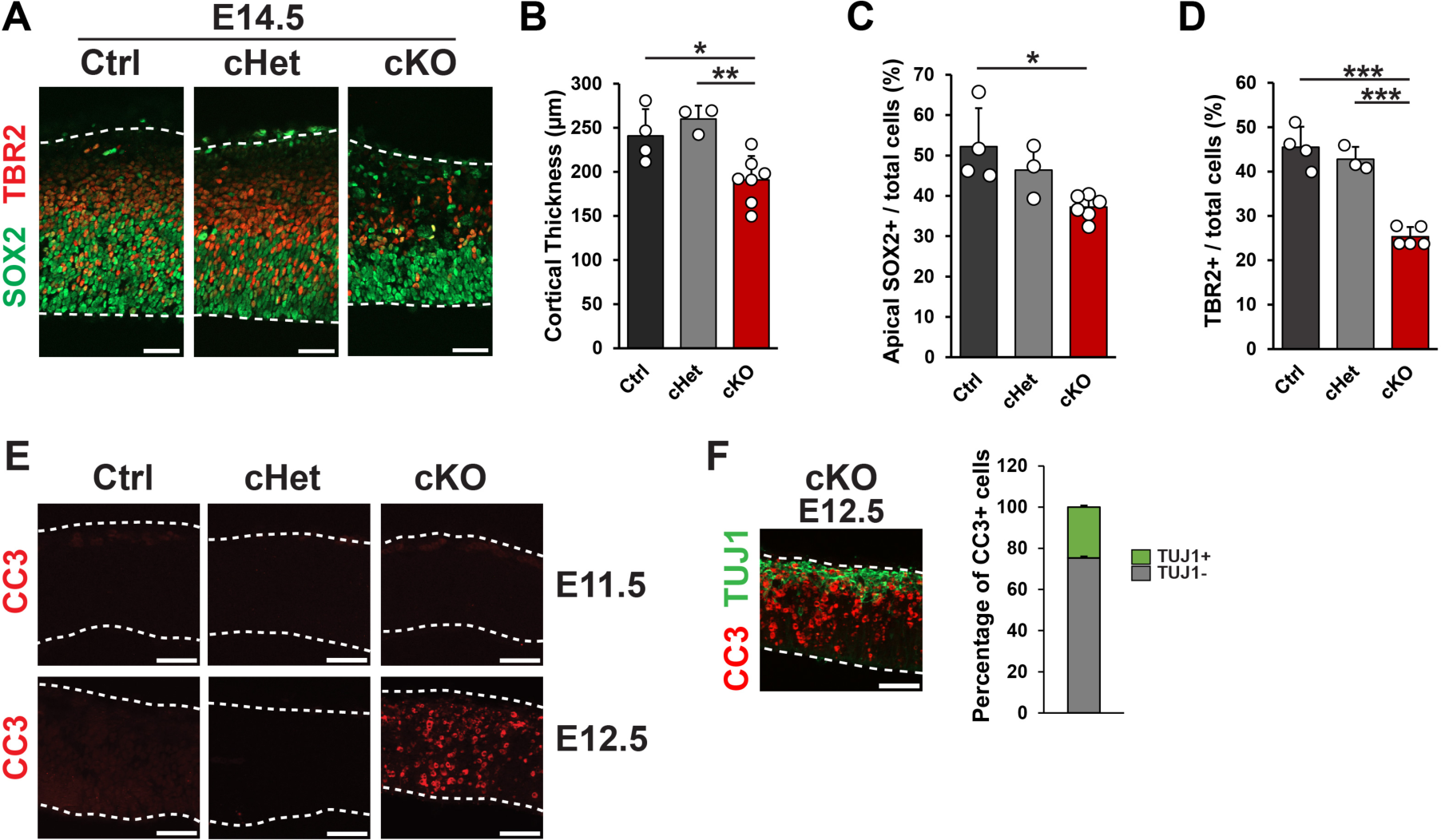
*Cnot3* loss impairs progenitor number and causes apoptosis. (A) Immunofluorescence for SOX2 and TBR2 in E14.5 cortical sections, with quantifications shown in (B-D). n = 3-6 embryos per genotype. (E) Immunofluorescence for CC3 in E11.5 (top) and E12.5 (bottom) cortical sections. (F) Immunofluorescence for CC3 and TUJ1 in E12.5 cortical sections, with quantification shown to the right (n = 3 embryos). \**p* < 0.05, \*\**p* < 0.01, \*\*\**p* < 0.001. One-way ANOVA with Tukey’s HSD post-hoc test. Error bars represent standard deviation. Scale bars: 50 µm.

The reduction in cortical thickness and decreased progenitor and neuronal cell density could be associated with alterations in proliferation or excessive cell death. We first assessed the latter possibility by measuring apoptosis. Staining for cleaved caspase 3 (CC3) revealed widespread cell death in cKO cortices beginning at E12.5 (Fig. 5E). Apoptotic cells were present throughout the cortex and were both TUJ1+ and TUJ1-. This indicates that *Cnot3* loss induces apoptosis of both newborn neurons and presumed progenitors (Fig. 5F).

We next sought to determine whether neuronal cell death was strictly a result of *Cnot3* loss in progenitors, or if *Cnot3* was also required in neurons for viability. For this we crossed *Cnot3*^lox/lox^ mice with *Nex*-Cre (Supplemental Fig. S4B), which is active in post-mitotic glutamatergic neurons beginning at E12.5 (Goebbels et al. 2006). Staining for CC3 at E18.5 indicated apoptosis in *Nex*-Cre cKO cortices (Supplemental Fig. S4C). This indicates that neuronal cell death is due, at least in part, to an intrinsic requirement of *Cnot3* for newborn neuron viability. Staining for TBR1, CTIP2, RORβ, and LHX2 at E18.5 revealed reductions of 33%, 20%, 18% and 10%, respectively, with only the deep layer neurons being significant (Supplemental Fig. S4D-H). The attenuated impact on superficial layers is most likely due to the later birthdate of these neurons and the time it takes for CNOT3 protein to be depleted after Cre-mediated recombination. Collectively, these data underscore the intrinsic requirement of CNOT3 for viability of multiple cell types of the developing cortex, including both neural progenitors and post-mitotic neurons.

### Rescue of apoptosis partially rescues *Cnot3* cKO phenotypes

Given that *Cnot3* is essential for cell viability, we next assessed the extent to which apoptosis contributes to the loss of both progenitors and neurons in *Emx1* cKO brains. Cell death in the cortex is often associated with upregulation of p53 signaling (Mao et al. 2016; Little and Dwyer 2019; Ulmke et al. 2021; Lupan et al. 2023). Consistent with this, we observed accumulation of p53 protein in E12.5 cKO cortices (Fig. 6A), indicating active p53 signaling at this stage. Given this, we generated *Emx1-*Cre;*Trp53^lox/lox^;Cnot3^lox/lox^*double cKO (dcKO) mice to abrogate p53-dependent apoptosis. Rescue of p53-dependent apoptosis was confirmed by the absence of p53 protein accumulation and CC3+ cells in E12.5 dcKO cortices (Supplemental Fig. S5A, B). Examination of dcKO brains at E18.5 revealed a striking rescue of cortical size (Fig. 6B, C). Notably, cortical thickness in dcKO brains was virtually unchanged compared to cKO (Fig. 6D). The density of early born, deep layer neurons marked by TBR1 and CTIP2 was similar between dcKO and cKO mice (Fig. 6E, F). In contrast, the density of later born neurons marked by RORβ and LHX2 was restored in dcKO mice (Fig. 6G, H). This suggests that p53-dependent apoptosis explains the loss of late-born, but not early-born neurons in *Cnot3* cKO mutants. As with density, the compound mutants also failed to rescue lamination patterns for deep layer markers, but did rescue lamination of superficial layers markers (Supplemental Fig. S5C-F). These data indicate both p53-dependent and p53-independent mechanisms for CNOT3 regulation of neuronal populations.

**Figure 6.**
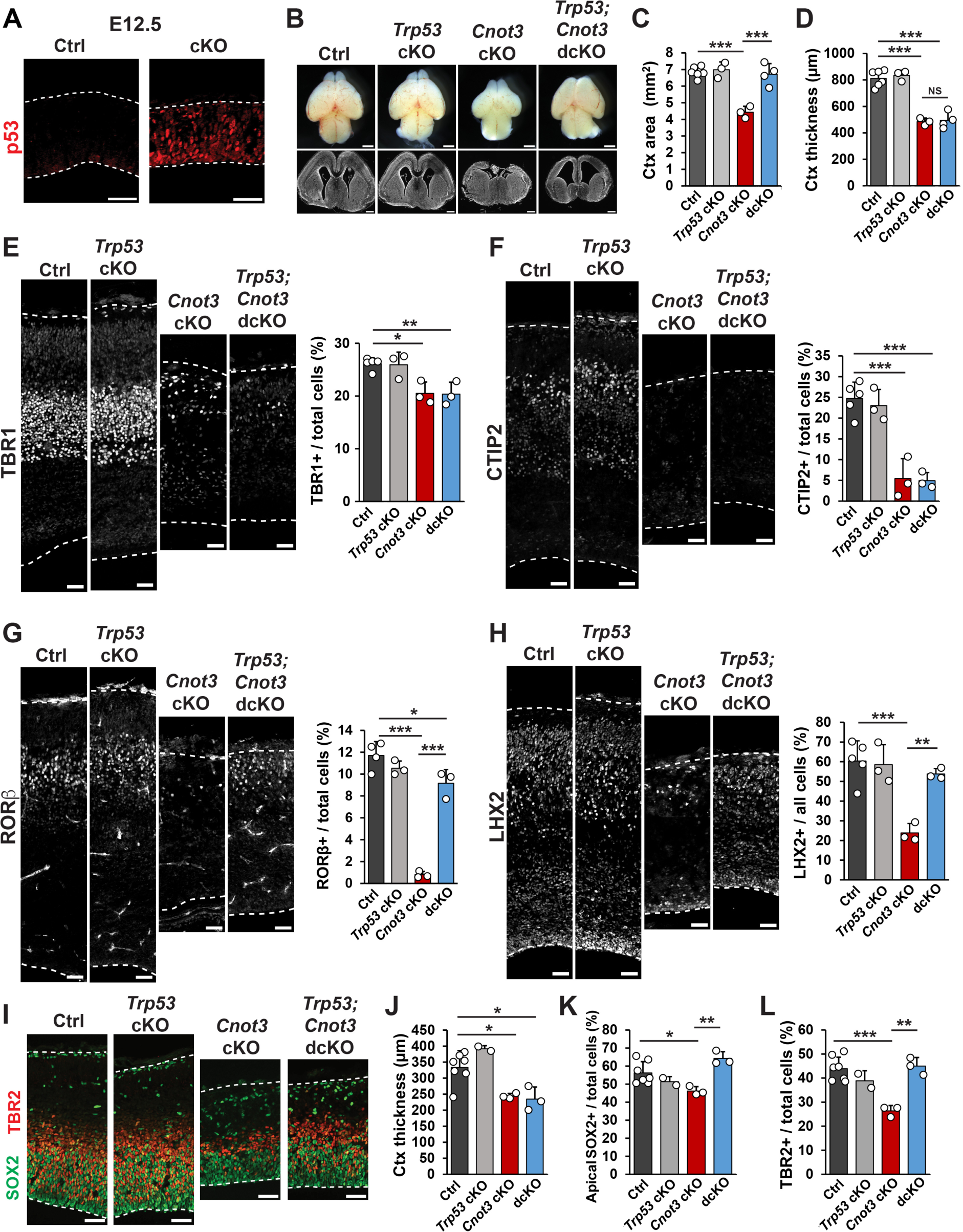
cKO of *Trp53* partially rescues *Cnot3* cKO phenotypes. (A) Immunofluorescence for p53 in E12.5 cortical sections. (B) Whole mount images and coronal cross sections of E18.5 brains. (C, D) Quantifications of cortex area and thickness (n = 3-6 embryos per genotype). (E-H) Immunofluorescence for indicated neuronal markers in E18.5 cortical sections, with quantifications shown to the right (n = 3-6 embryos per genotype). (I) Immunofluorescence for SOX2 and TBR2 in E14.5 cortical sections. (J-L) Quantification of cortical thickness, SOX2, and TBR2 at E14.5 (n = 2-7 embryos per genotype). \**p* < 0.05, \*\**p* < 0.01, \*\*\**p* < 0.001. One-way ANOVA with Tukey’s HSD post-hoc test. Error bars represent standard deviation. Scale bars: 1 mm (B, top), 500 µm (B, bottom), 50 µm (A, E-I).

The reduction in superficial layer neurons may be due to apoptosis of progenitors at stages when these neurons are being produced. To probe this possibility, we quantified neurogenesis of compound mutant brains at E14.5. Similar to E18.5, cortical thickness at E14.5 was unchanged in dcKO brains compared to cKO (Fig. 6I, J). However, the densities of apical SOX2+ RGCs and TBR2+ IPs were rescued in dcKO brains (Fig. 6K, L). Thus, the decrease in progenitor density in E14.5 *Emx1-*cKO brains is due to p53-dependent apoptosis. This observation is consistent with the rescue of upper layer neurons, whose peak production occurs around E14.5. Collectively, our analysis of dcKO mice suggests that *Cnot3-*dependent microcephaly and neuronal loss are due in part to massive apoptosis. Notably, p53-dependent apoptosis is developmentally specific as the loss of upper layer neurons is p53-dependent whereas loss of deep layer neurons is largely p53-independent.

### CNOT3 is required for regulation of cell cycle duration and cell fate

Given the findings above, we next asked if *Cnot3* controls the rate at which early-stage progenitors produce neurons. For this, we employed a previously established *in vivo* semi-cumulative labeling strategy to measure total cell cycle (Tc) and S-phase (Ts) durations at E12.5 (Quinn et al. 2007). BrdU was intraperitoneally injected into pregnant dams to label DNA in S-phase progenitors, followed by a second injection with EdU 1.5 hr later (Fig. 7A). Embryonic brains were dissected at t = 2 hr and stained for BrdU, EdU, and Ki67 (proliferative cells). Total cell cycle and S-phase in control cortices were determined to be ∼10 hr and ∼5 hr, respectively (Fig. 7B). Importantly, both measurements are in close agreement with previous measurements (Takahashi, Nowakowski and Caviness 1995; Boyd et al. 2015). In contrast, in cKO cortices, total cell cycle and S-phase durations were both significantly increased to ∼14 hr and ∼6.5 hr, respectively (Fig. 7B). The ratio of Ts/Tc was unchanged across the three genotypes, indicating that cell cycle lengthening was not biased towards S-phase (Fig. 7C). These data demonstrate that *Cnot3* loss from early-stage progenitors is associated with overall slower cell cycle.

**Figure 7.**
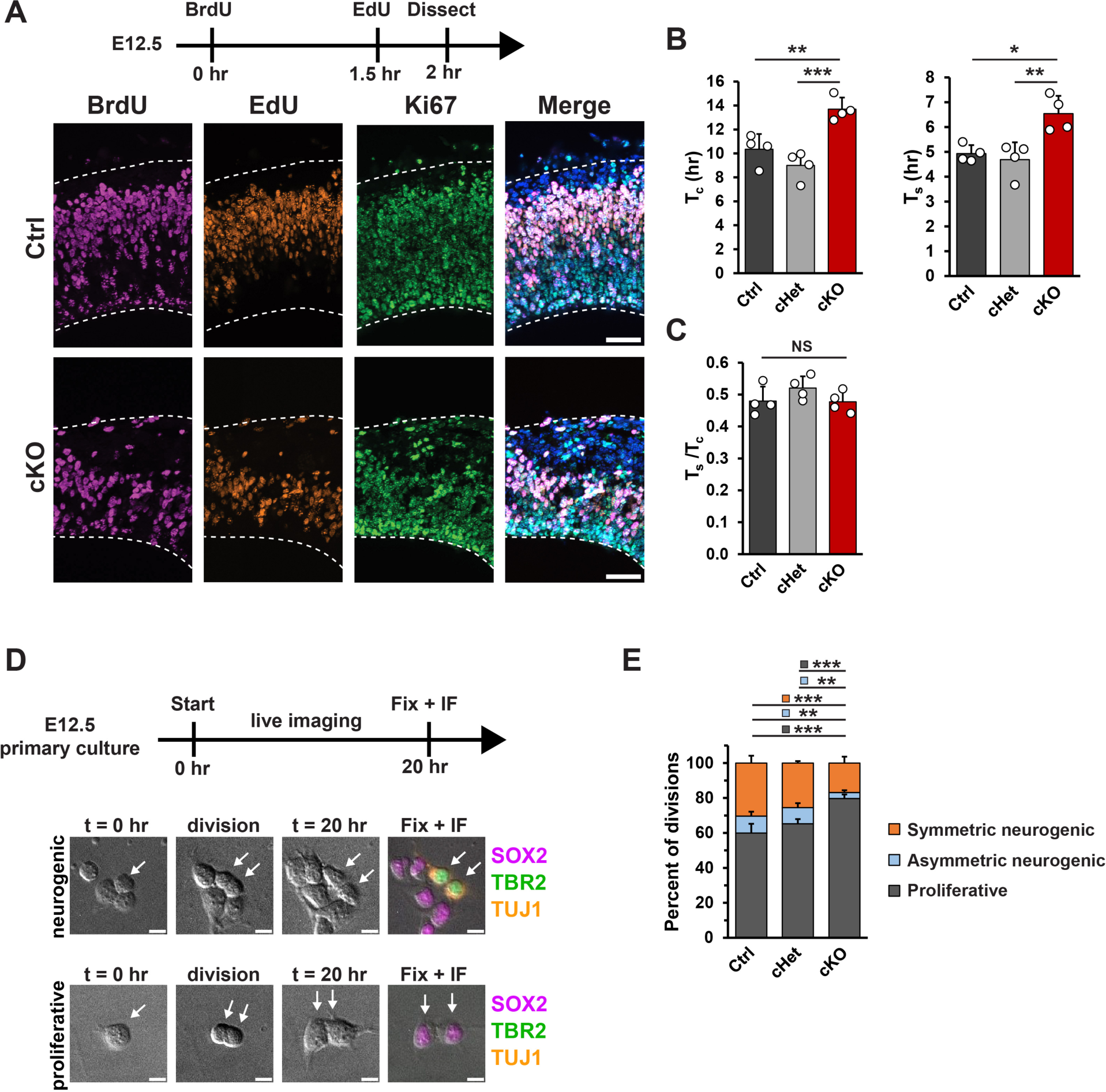
*Cnot3* is required for progenitor cell cycle duration and neurogenic divisions. (A) Top, schematic of experimental approach for semi-cumulative labeling at E12.5. Bottom, immunofluorescence for the indicated markers in E12.5 cortical sections. (B) Quantification of total cell cycle duration (T_c_), S-phase (T_s_) (n = 4 embryos per genotype). (C) Quantification of the ratio between S-phase and total cell cycle (T_s_/T_c_). (D) Top, schematic of experimental approach for live imaging of E12.5 primary progenitor divisions. Bottom, DIC images at the beginning (t = 0 hr) and end (t = 20 hr) of the live imaging session and immunofluorescence after fixation for SOX2, TBR2, and TUJ1 showing representative proliferative and neurogenic divisions (n: Ctrl = 4 brains (446 divisions), cHet = 3 brains (331 divisions), cKO = 3 brains (350 divisions)). (E) Quantification of cell divisions: Proliferative (2 SOX2+TUJ1-, 2TBR2+TUJ1-, or one of each); Asymmetric neurogenic (1 SOX2+TUJ1- or TBR2+TUJ1- and 1 TUJ1+); Symmetric neurogenic (2 TUJ1+). \**p* < 0.05, \*\**p* < 0.01, \*\*\**p* < 0.001. One-way ANOVA with Tukey’s HSD post-hoc test (B, C). Chi-squared test with Bonferroni post-hoc adjusted *p*-values (E). Error bars represent standard deviation. Scale bars: 50 µm (A), 10 µm (D).

We next investigated the extent to which *Cnot3* controls early stage progenitors’ ability to produce neurons. For this we performed a live imaging assay to monitor the fate of dividing progenitors and their progeny (Fig. 7D). Primary cell cultures were prepared from E12.5 control, cHet, and cKO cortices, and progenitors were imaged every 10 min for a period of 20 hr, as previously (Pilaz et al. 2016; Hoye et al. 2022). After 20 hr, cells were fixed and stained for SOX2 (RGCs), TBR2 (IPs), and TUJ1 (neurons) to assign the fate of progeny. In control progenitors, ∼60% of divisions were proliferative (two SOX2+ or TBR2+ progeny), ∼10% were asymmetric neurogenic (one SOX2+ or TBR2+, and one TUJ1+ progeny), and ∼30% were symmetric neurogenic (two TUJ1+ progeny). In contrast, divisions of cKO progenitors were skewed towards producing more progenitors, with ∼80%, 5% and 15% of divisions being proliferative, asymmetric neurogenic, and symmetric neurogenic, respectively (Fig. 7E). cHet progenitors had an intermediate phenotype that was not statistically significant. The observation that progenitors undergo fewer neurogenic divisions in cKO cells suggests that *Cnot3* loss from progenitors directly impairs neuronal production. This, together with prolonged cell cycle, suggest potential mechanisms to help explain the p53-independent loss of deep layer neurons.

### Upregulation and impaired turnover of non-optimal mRNAs in cKO cortices

Next, we aimed to understand the molecular consequences of *Cnot3* loss in the developing cortex. For this, we isolated RNA from E12.5 control, *Cnot3* cKO, and *Cnot3;Trp53* dcKO cortices and performed bulk RNA-seq. We chose E12.5, the earliest onset of phenotypes in cKO embryos, with the goal of understanding primary molecular deficits. Using cutoffs of *p_adj_* < 0.05 and fold-change > 2, we identified 1,216 transcripts that were differentially expressed in cKO cortices compared to control, and 1,164 differentially expressed transcripts in dcKO cortices (Fig. 8A, and Supplemental Table S5). As expected, p53 target genes, including *Sesn2, Eda2r, Ccng1,* and *Cdkn1a*, were among the most upregulated genes in cKO cortices (Fig. 8A, B). In dcKO cortices, these transcripts were each expressed at levels similar to control, confirming attenuation of p53-dependent apoptosis in the compound mutant mice. These gene expression changes were each independently validated by RT-qPCR (Fig. 8B). This further supports p53-dependent apoptosis as a major mechanism of *Cnot3* cKO phenotypes. Additionally, multiple transcripts involved in cell cycle regulation, including *Ccnd1, Ccnd2,* and *Cdk4*, were downregulated in either cKO or dcKO cortices (Supplemental Fig. S6A), providing molecular support for cell cycle phenotypes reported above.

**Figure 8.**
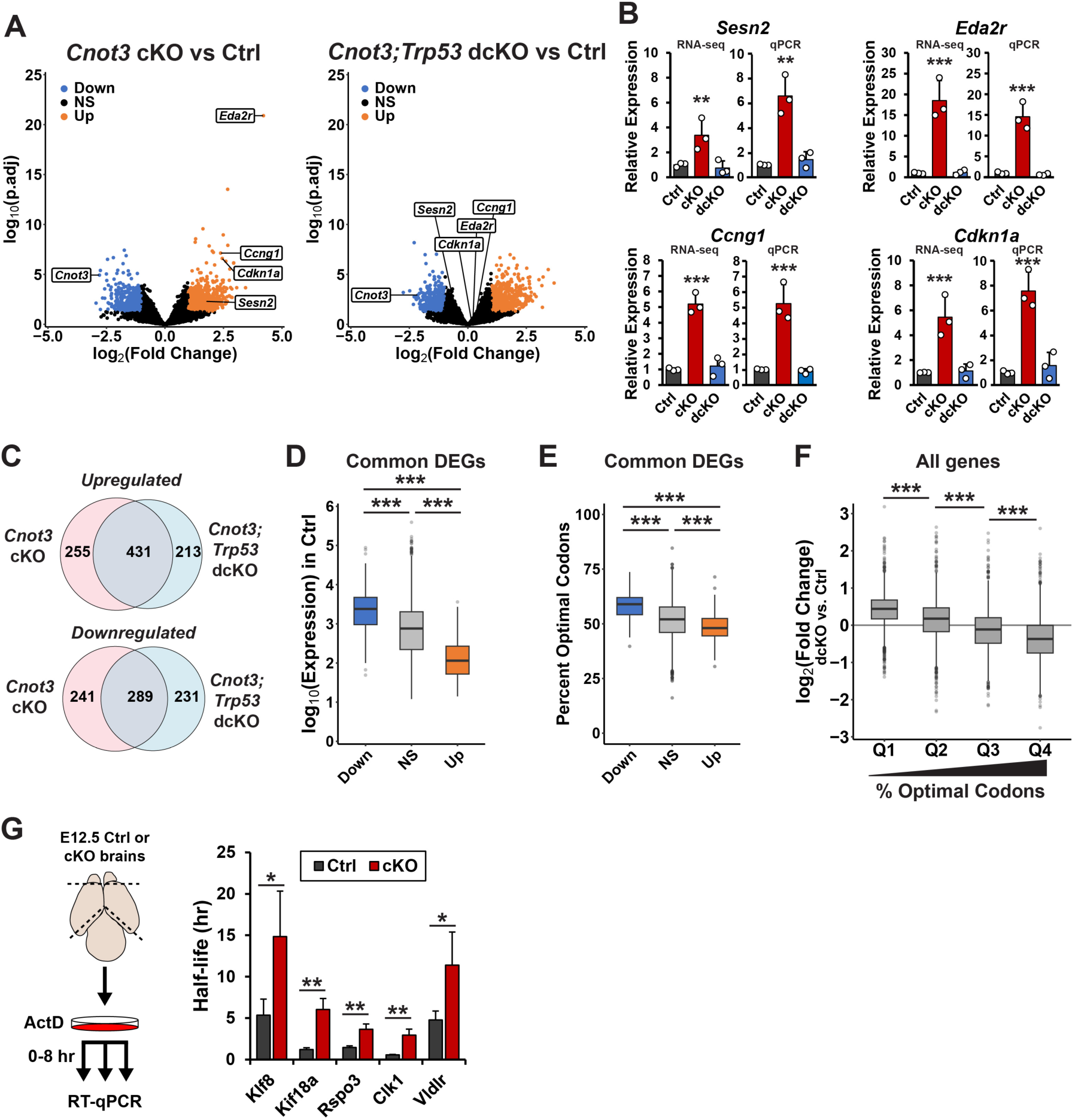
Upregulation and impaired turnover of non-optimal mRNAs in *Cnot3* cKO cortices. (A) Volcano plots showing differentially expressed transcripts in cKO (left) and dcKO (right) cortices versus control at E12.5 (n = 3 embryos per genotype). (B) Expression of p53 target genes as measured by RNA-seq (left) or qPCR (right). (C) Venn diagram showing overlap between differentially expressed genes in cKO and dcKO cortices. (D) Log_10_(expression) of commonly upregulated genes (Up; n = 431), commonly downregulated genes (Down, n = 289), and all other genes (NS, n = 13,089) in Ctrl cortices. (E) Percentage of optimal codons (CSC > 0) for commonly differentially expressed genes. (F) Log_2_(fold change) in dcKO versus control cortices for all genes, binned into quartiles based on percentage of optimal codons. (G) Left, schematic of experimental approach for transcriptional shutoff of E12.5 primary cultures with actinomycin D (Act D). Right, half-lives of the indicated transcripts determined by RT-qPCR (n = 4 embryos per genotype). \**p* < 0.05, \*\**p* < 0.01, \*\*\**p* < 0.001. Adjusted *p*-values from DESeq2 (B (RNA-seq)). One-way ANOVA with Tukey’s HSD post-hoc test (B (qPCR), D-H), Welch’s two-sample t-test (G). Error bars represent standard deviation.

Among the 686 transcripts that were upregulated in cKO cortices, 431 (63%) were also upregulated in dcKO cortices (Fig. 8C). A similar overlap was observed for downregulated transcripts, with 289 in common out of the 530 downregulated transcripts in cKO cortices (55%). We focused our subsequent analysis on these commonly differentially expressed transcripts as they are most likely to be regulated by *Cnot3,* independent of apoptosis. GO analysis of differentially expressed transcripts revealed terms related to cell projections and neuronal development (Supp. Fig. 6B, C).

To further define the molecular features of these shared *Cnot3*-dependent transcripts, we assessed their baseline expression levels in control cortices. Upregulated transcripts were expressed at significantly lower levels compared to *Cnot3*-independent transcripts (Fig. 8D). In comparison, downregulated transcripts had significantly higher levels of expression in control brains (Fig. 8D). These data suggest that *Cnot3* plays a role in maintaining correct gene expression levels in cortical cells.

Previous work has shown that the CCR4-NOT complex is recruited to mRNAs with low codon optimality in a manner that is dependent on CNOT3 interaction with the ribosome E-site, and deletion of the yeast homolog of CNOT3 (Not5) preferentially stabilizes non-optimal mRNAs (Buschauer et al. 2020; Absmeier et al. 2023). We thus evaluated the relationship between expression changes and codon content using CSC values calculated from our SLAM-seq data in WT cells (Fig. 1H). Comparing optimal codon content among differentially expressed transcripts showed that upregulated transcripts were significantly less optimal than unchanged transcripts, while downregulated transcripts were more optimal (Fig. 8E). We probed the relationship between codon content and expression changes in mutant cortices further by binning all mRNAs into quartiles based on percentage of optimal codons. This analysis revealed that the most non-optimal mRNAs tended to be the most upregulated in *Cnot3* mutants (Fig. 8F). Collectively, these data implicate *Cnot3* in regulating RNA expression in a manner dependent on codon content, with preferential upregulation of non-optimal mRNAs in mutant cortices. This further supports RNA stability control as a mechanism by which *Cnot3* regulates cortical development.

Disruption of the CCR4-NOT complex in *Cnot3* cKO cortices is expected to stabilize target transcripts by impairing deadenylation. To evaluate whether increased expression of non-optimal mRNAs is due to changes in mRNA stability, we measured mRNA half-lives in primary cortical cells using transcriptional shutoff with Actinomycin D (ActD). Primary cultures were generated from E12.5 control and cKO cortices and cells were harvested 0, 2, and 8 hr after addition of ActD to assess RNA degradation by RT-qPCR (Fig. 8G). We examined the decay kinetics of five mRNAs whose expression increased in cKO cortices (Supplemental Fig. S6D) and that fell into the lowest quartile of codon optimality. All five transcripts were stabilized in cKO cells, consistent with defective mRNA turnover in the absence of *Cnot3* (Fig. 8G and Supplemental Fig. S6E). Of note, three of these genes (*Klf8, Kif18a,* and *Clk1*) have roles in cell cycle regulation (Zhao et al. 2003; Stumpff et al. 2008; Dominguez et al. 2016). A fourth gene (*Vldlr*) is a reelin receptor important for cortical neuron positioning (Hack et al. 2007). In sum, these data reveal upregulation of non-optimal, poorly expressed RNAs in cKO cortices. Transcriptional shut-off analysis of select transcripts confirms impaired RNA degradation in the absence of *Cnot3*. These observations directly implicate the CCR4-NOT complex in controlling gene expression in the developing cortex.

## Discussion

RNA regulation in the developing cortex is a dynamic process, with rapid changes in gene expression occurring across dual axes of time and differentiation. RNA expression levels are determined by complementary rates of synthesis and degradation, but the quantitative contribution of the latter to cortical development has been largely unknown. Here, we used molecular and genetic approaches to understand how RNA turnover controls cortical development. Our transcriptome-wide survey of RNA half-lives across development provide an in-depth understanding of the *cis* factors that contribute to RNA turnover, as well as previously unappreciated relationships between RNA stability and developmental gene expression changes. We further use genetic manipulation of the CCR4-NOT complex to demonstrate the consequences of misregulating RNA turnover *in vivo*, finding that CNOT3 is critical for neurogenesis. Our findings reinforce essential roles for RNA stability regulation in cortical development, highlighting new mechanisms relevant for related neuropathologies.

### A new landscape of mRNA stability in the developing cortex

Our study reveals new post-transcriptional regulatory layers which balance transcriptional control to collectively govern gene expression in the developing cortex. We discover a relationship between developmental expression changes and half-life. Specifically, we find that transcripts that increase in expression at later stages of cortical development tend to have longer half-lives, while transcripts that decrease in expression tend to have shorter half-lives (Fig. 9A). This demonstrates that RNA stability is tied to dynamic gene expression over the course of development.

**Figure 9.**
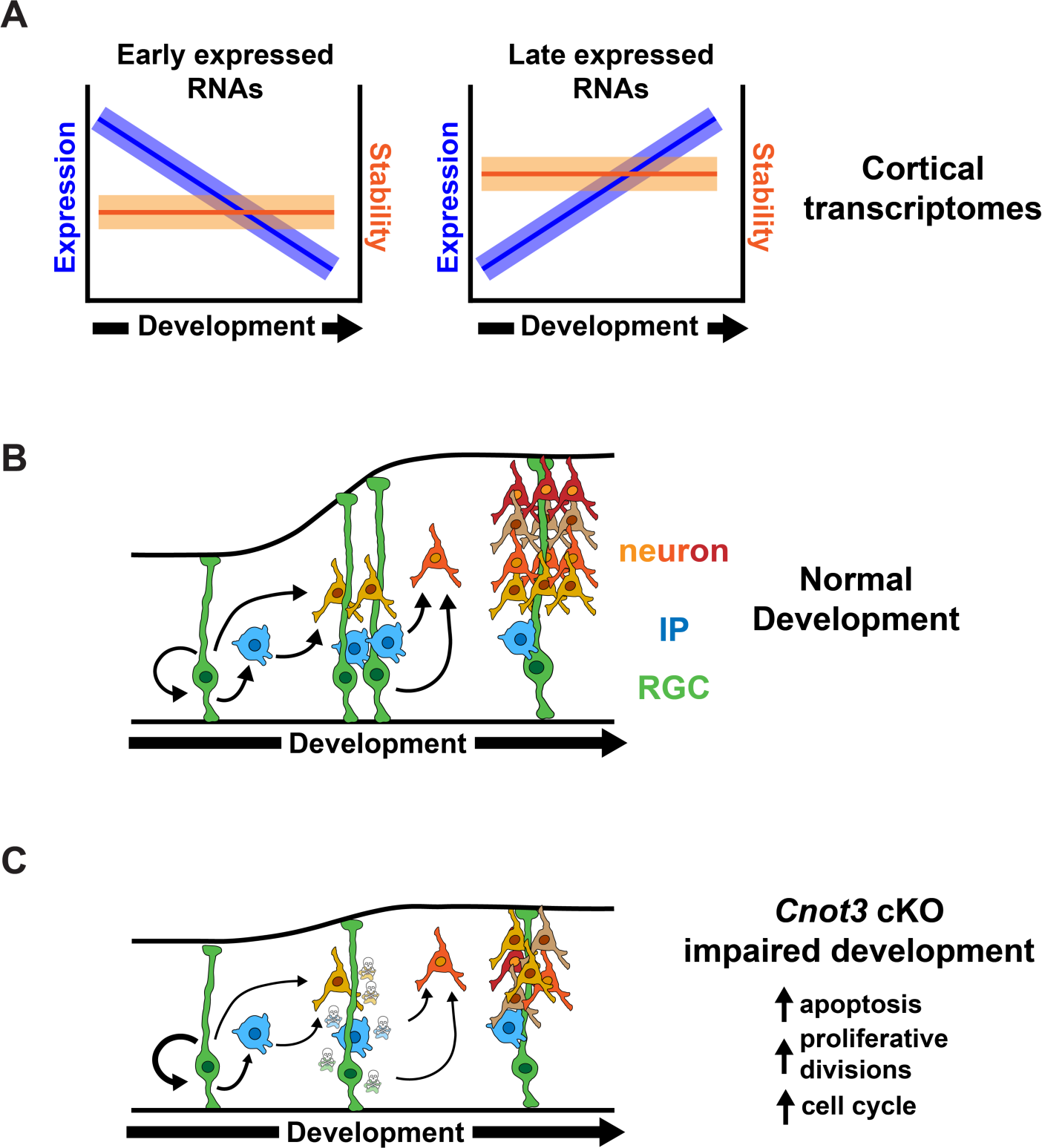
Models for relationship between RNA turnover and gene expression, and the role of *Cnot3* in cortical development. (A) Transcripts that are downregulated across development are relatively unstable, while those that are upregulated across development are relatively stable. For the majority of transcripts, RNA half-lives are static across development. Cortical development in WT (B) and *Cnot3* cKO embryos (C). Loss of *Cnot3* impacts development by p53-dependent and p53-independent mechanisms leading to apoptosis of progenitors and neurons, increased proliferative divisions, and longer cell cycle duration.

Our data help elucidate the extent to which RNA degradation varies across developmental time and space. We provide evidence for two distinct modes of RNA stability regulation in the cortex. First, a dynamic half-life model in which the stability of a given RNA varies across development. Second, a static half-life model in which RNA stability depends on when it is most highly expressed during development. In this latter paradigm, RNAs only required early in development have short half-lives to promote their rapid clearance. In contrast, RNAs needed later in development are relatively more stable to promote their accumulation over time. While these models are not mutually exclusive, our data indicates that the static half-life model is predominant in the developing cortex, as RNA decay kinetics for most transcripts are unchanged across development. While dynamic half-life regulation occurs for a subset of transcripts (Fig. 2), this appears to be the exception rather than the rule.

Our experimental design allowed us to assess RNA half-lives as cellular composition of the cortex transitions from progenitor-enriched at E11.5 to neuron-enriched at E16.5. If the half-life of a given transcript greatly differed between progenitors and neurons, we expect to have observed a shift in RNA stability kinetics from E11.5 to E16.5. However, we did not observe widespread changes in half-lives across these developmental transitions. This suggests that turnover rates for most transcripts may be similar regardless of the cell type in which they are expressed. While we used scRNA-seq datasets and knowledge of cell composition changes from E11.5 to E16.5 to inform cell-specific RNA stability, our study did not directly compare RNA decay in progenitors and neurons. Strategies for single cell analysis of RNA stability have successfully been employed in complex model systems including intestinal organoids and zebrafish embryos (Battich et al. 2020; Fishman et al. 2024). Adapting these techniques for *in vivo* analysis of RNA stability across diverse cell types of the developing mammalian cortex is an exciting direction for future investigation.

### The CCR4-NOT complex is essential for cortical development

Our study used an orthogonal genetic approach to demonstrate essential roles of RNA stability in cortical development by focusing on the CCR4-NOT complex. To that end, we used progenitor and neuronal Cre drivers to assess requirements of *Cnot3* in neural progenitors and newborn neurons of the developing cortex. While we specifically targeted CNOT3 in progenitors, expression of the central scaffold CNOT1 was also affected, suggesting the entire complex may be affected (Fig. 4C). *Cnot3* cKO brains are microcephalic, with fewer progenitors and neurons. We show these phenotypes are due in part to widespread p53-dependent apoptosis of both populations. Interestingly, *Cnot3* cKO does not induce apoptosis in mouse ESCs or male germ cells (Zheng et al. 2016; Chen et al. 2023), but does in B cells by directly regulating the stability of *p53* mRNA(Inoue et al. 2015). These observations collectively suggest that the requirement of *Cnot3* for mammalian cell viability may be cell type specific.

Our experiments using compound mutants revealed that *Cnot3* controls cortical neurogenesis via both p53-dependent and p53-independent mechanisms (Fig. 9B, C). By analyzing cKO and dcKO brains at E14.5 and E18.5 we defined the timing of when these mechanisms are predominant. We observed a preferential rescue of superficial layer neuron density at E18.5, coupled with the rescue of progenitor density at E14.5, when superficial neurons are being born. This suggests that p53-dependent mechanisms are especially prevalent at later stages of cortical neurogenesis. In contrast, loss of early born deep layer neurons was largely p53-independent.

This raises the question of what mechanisms are at play at earlier developmental stages? We determined that E12.5 progenitors take longer to divide and undergo more proliferative divisions at the expense of neurogenic divisions. These progenitor defects are ultimately expected to produce less neurons across development. We predict that this change in cell fate contributes to genesis of fewer deep Layer VI and Layer V neurons (TBR1+ and CTIP2+) in dcKO cortices. How *Cnot3* controls progenitor cell cycle and neurogenic divisions is a fascinating question for future studies. It is notable that similar phenotypes have been observed for mutations in other RNA binding proteins (Hoye and Silver 2021), reinforcing the role of post-transcriptional control in cell fate specification.

### CNOT3 as a regulator of gene expression in the developing cortex

To understand the molecular underpinnings of these cortical development phenotypes, we performed RNA-seq of control, cKO, and dcKO cortices. This allowed us to measure transcriptomic changes due to p53-dependent cell death versus those which are p53-independent. Consistent with p53-dependent mechanisms, we observed increased expression of p53 transcriptional targets in cKO cortices, including *Eda2r, Sesn2,* and *Ccng1*. We also observed decreased expression of several cell cycle related transcripts, including *Ccnd1, Ccnd2,* and *Cdk4*. These observations align with our *in vivo* measurements of prolonged cell cycle, as well as previous reports indicating a role for CNOT3 in cell cycle regulation (Takahashi et al. 2012; Zhou et al. 2017; Shirai et al. 2019). Of note, *Cdk4* is also downregulated upon knockdown of *CNOT3* in human cardiomyocytes, suggesting a conserved mechanism of cell cycle regulation.

The CCR4-NOT complex is primarily described as a regulator of mRNA stability, however it has roles in multiple aspects of RNA metabolism (Collart 2016). Furthermore, shortening of poly(A) tails not only impacts RNA stability, but can also contribute to changes in translation efficiency (Passmore and Coller 2022). While we cannot rule out the possibility that cKO phenotypes are caused in part by disruption of other molecular processes, several lines of evidence suggest that RNA turnover is impaired in cKO cortices. First, gene expression changes in cKO and dcKO cortices correlated with codon content, with non-optimal mRNAs showing the highest upregulation. In light of previously established roles of the CCR4-NOT complex in degrading non-optimal RNAs (Webster et al. 2018; Buschauer et al. 2020), this is consistent with widespread RNA stabilization in mutant cortices. We confirmed this hypothesis using transcriptional shutoffs to directly show stabilization of non-optimal mRNAs in cKO cells. Overall, this data highlights the importance of maintaining control of RNA degradation in the developing cortex.

The CCR4-NOT complex, including CNOT3, is increasingly implicated in human disease, such as brain malformations and autism (Kruszka et al. 2019; Martin et al. 2019; Meyer et al. 2020; Vissers et al. 2020; Priolo et al. 2021; Lv et al. 2023; Zhao et al. 2023). This highlights the need to understand the functions of this complex in the brain and how its molecular perturbation causes human disease. Importantly, this work establishes an essential role for CNOT3 in cortical development and lays the groundwork for future studies to directly investigate how disease-causing mutations impact the developing cortex.

## Materials and Methods

### Mice

Animal use was approved by the Duke Institutional Animal Care and Use Committee and followed ethical guidelines provided by the Duke Division of Laboratory Animal Resources. Mouse lines were previously described: *Emx1*-Cre JAX stock #005628 (Gorski et al. 2002), *Trp53*^lox/lox^ JAX stock #008462 (Marino et al. 2000), *Nex*-Cre (Goebbels et al. 2006), *Cnot3^lox/lox^*(Zheng et al. 2016). Primers used for genotyping are listed in Supplemental Table S6. For embryo staging, plug dates were defined as embryonic day (E)0.5 on the morning the plug was identified.

### SLAM-seq

#### Sample collection and library preparation

Embryos were dissected in cold PBS and cortical tissue was microdissected from three biological replicates. Each biological replicate consisted of pooled cortical tissue from 7 embryos (E11.5), or 2 embryos (E14.5 and E16.5). Cortices were dissociated by incubation with 0.25% Trypsin (Thermo Fisher) at 37 °C for 5 min (E11.5), 10 min (E14.5) or 20 min (E16.5). Trypsinization was halted by addition of trypsin inhibitor (Sigma T6522) diluted in neural progenitor media [DMEM (Thermo Fisher 11995-065) supplemented with N-acetyl-L-cysteine (Sigma A9165, 1 mM), N2 (Thermo Fisher 17502048, 1X), B27 without vitamin A (Thermo Fisher 12587-010, 1X), and mouse bFGF (R&D Systems 3139-FB025, 10 ng/mL)]. Following trituration with a P1000 pipette, cells were pelleted at 200 g for 5 min to remove debris then resuspended in fresh neural progenitor media. Cells from each biological replicate were divided into five wells of a 24-well plate coated with poly-D-lysine (Sigma P7280) and placed at 37 °C in a humidified incubator with 5% CO_2_. Cells were allowed to attach to the plate for 2 hr, then media was replaced with fresh neural progenitor media containing 100 µM 4sU (Sigma T4509). The time of 4sU addition was designated as the t = -20 hr time point. Media was replaced with fresh 4sU containing media at t = -12, -8, -4, and -1 hr. At t = 0 hr, a 50X molar excess (i.e. 5 mM) of uridine was added to the media. Cells from a single well per replicate were lysed 0, 2, 4, 8, and 12 hr after addition of uridine by addition of RLT buffer (Qiagen RNeasy Plus Micro Kit, cat. 74034) supplemented with 1 mM DTT. Lysates were stored at -80 °C before further processing. RNA was extracted from lysates using the Qiagen RNeasy Plus Micro Kit according to manufacturer’s instructions, except all buffers were supplemented with 1 mM DTT. Alkylation of 4sU RNA was performed as previously described (Herzog et al. 2017) using ∼1.5 µg of RNA per replicate. After alkylation, RNA was re-purified using the RNA Clean and Concentrator Kit (Zymo 11-325). Sequencing libraries were generated using 400 ng of alkylated input RNA per sample and the Illumina Stranded mRNA Prep Kit (cat. 20040532) according to the manufacturer’s instructions.

#### Read mapping and half-life calculation

Sequenced reads were processed with the SlamDunk tool v0.4.3, using the mm10 genome and exons extracted from the GENCODE vM24 gene annotation. The number of covered (T)s and converted (T)s for each exon was extracted from the SlamDunk results. Covered (T) counts for all exons within a gene were summed together, and the same was done for converted (T)s. A conversion rate was calculated for each gene as the percentage of converted (T)s to covered (T)s. These conversion rates were calculated for each time point, and a half-life value was calculated for each gene based on the decay in these conversion rates over time. The decay curve was estimated in R with the “nlsLM” function from "minpack.lm" package, using the model “y ∼ exp(-(k*x))”, where “x” are the time points and “y” are the conversion rates scaled such that the initial conversion rate (at time point 0) is 1. The “k” value was extracted from the fit curve, and the half-life was calculated as “ln(0.5) / -k”.

#### Analysis

##### Quality control

After calculating half-lives, the data was filtered for read coverage and the quality of half-life calculation. The following filters were applied to each transcript within each biological replicate: coverage on T > 500 at each time point of the chase, half-life < 24 hr and > 0 hr, and R^2^ > 0.6. Transcripts passing these filtering steps in all three biological replicates per stage were retained for further analysis and are listed in Supplemental Table S1.

##### GO Analysis

Gene ontology analysis for top 10% most and least stable transcripts at each stage were performed using Panther (Bankhead et al. 2017).

##### Codon stabilization coefficient (CSC)

Mouse coding region sequences for were obtained from Ensembl using the biomaRt package in R (Durinck et al. 2005). For genes with multiple transcript isoforms, the longest isoform was used. Codon frequency for each transcript was calculated using the oligonucleotideFrequency function within the Biostrings(Lifschitz et al. 2022) package using width = 3, step = 3, and as.prob = T. CSC for each codon was calculated as the Pearson correlation coefficient between codon frequency and transcript half-life. Optimal codons were defined as those with CSC > 0, and non-optimal codons were defined as those with CSC < 0.

##### maSigPro

For statistical analysis of half-life changes across each of the three stages, transcript lists were filtered to include only those that were present in all three data sets. Half-lives were z-score normalized and transcripts with significant differences across all three stages were identified using the maSigPro(Nueda, Tarazona and Conesa 2014) package in R. The following modifications were made from the default parameters: for the regression fitting T.fit function, alfa = 0.05 was used, to extract the significant genes using the get.siggenes function, rsq = 0.6 was used. Significant genes were then clustered using the pheatmap function in R with clustering_method = “ward.D2”.

##### Differential Expression

Differential expression between stages was determined using DESeq2 (ref. (Love, Huber and Anders 2014)). Count data from the t = 0 hr time point for each replicate was used to generate the input count matrix. Data was filtered to include only those genes that had >10 reads in all samples. Significantly differentially expressed genes between E11.5 and E14.5, or E14.5 and E16.5 were defined as those with |fold-change| > 2 and *p_adj_* < 0.05.

##### Generating progenitor and neuron enriched gene lists

Lists of genes enriched in progenitors or neurons were generated using two previously published scRNA-seq datasets (Telley et al. 2019; Di Bella et al. 2021). Independent enrichment lists were generated from each dataset, and genes that were in common between each were used in this study (Supplemental Table S4). For Telley et al., data was accessed online (http://genebrowser.unige.ch/telagirdon/) and the “Genes with similar dynamics” feature was used to generate three lists of the top 100 genes most similar to *Sox2* (RGCs), *Eomes* (IPs), and *Neurod2* (neurons). The *Sox2* and *Eomes* lists were combined to create the progenitor list for the Telley et al. dataset. For Di Bella et al., the gene expression matrix was downloaded from the Single Cell Portal (https://singlecell.broadinstitute.org/single_cell/study/SCP1290/molecular-logic-of-cellular-diversification-in-the-mammalian-cerebral-cortex). The FindMarkers function in Seurat(Hao et al. 2024) was used to identify genes that are at least 1.5 fold enriched and expressed in at least 50% of cells of the desired cell type. Genes found to be enriched in “Apical Progenitors” and “Intermediate Progenitors” were combined to generate the progenitor-enriched list. Genes enriched in “Immature Neurons”, “DL CPN”, and “UL CPN” were combined to generate the neuron-enriched list.

### Western blots

E12.5 cortices were homogenized in cold RIPA buffer supplemented with Halt Protease Inhibitor (Thermo Fisher 7834). Lysates were incubated on ice for 10 min then clarified by centrifugation at 15,000 rpm for 5 min at 4 °C. For western blot, 10 µg of protein lysate was mixed with Laemmli buffer (1x final) and DTT (25 mM final) and boiled at 95 °C for 5 min. Samples were loaded into Mini-PROTEAN TGX precast gels (Bio-Rad) and run at 100-150V for 1-2 hr. Transfer was performed using the Trans-Blot Turbo Transfer System (Bio-Rad). Membranes were blocked in 5% milk in TBST then incubated with primary antibodies overnight at 4 °C. The following primary antibodies were used: CNOT1 (CST 44613S, 1:1000), CNOT2 (Thermo Fisher 10313-1-AP, 1:1000), CNOT3 (Thermo Fisher 11135-1-AP, 1:500), ACTB (Santa Cruz sc4778, 1:500). After primary antibody incubation, blots were washed three times with TBST, then incubated with secondary antibody for 1 hr at RT. The following secondary antibodies were used: goat-anti-mouse-HRP (1:5000), goat-anti-rabbit-HRP (1:5000). After secondary antibody incubation, blots were washed three times with TBST, then exposed to Pierce ECL Western Blotting Substrate (Thermo Fisher). Imaging was performed using the Gel Doc XR system (Bio-Rad). Quantification was performed using ImageJ.

### Immunofluorescence

Embryonic brains were fixed in 4% PFA at 4 °C overnight, washed 3 x 10 min in cold PBS, then incubated in 30% sucrose at 4 °C overnight. Brains were frozen in NEG-50 (Fisher), and 20 µm thick coronal sections were collected on charged glass slides and stored at -80 °C until staining. For IF, frozen sections from the somatosensory cortex were thawed for 5-10 min at RT then washed twice in PBS. For RORβ staining only, antigen retrieval was performed by boiling sections in sodium citrate buffer (10 mM sodium citrate, 0.05% Tween 20, pH 6.0) for 20 min, then cooling to RT for 10 min followed by three PBS washes. Sections were permeabilized with 0.3% TritonX-100 in PBS for 15 min and blocked in 5% NGS in PBS for 1 hr at RT. Sections were incubated with primary antibodies for 2 hr at RT and secondary antibodies for 30 min at RT. The following primary antibodies were used: SOX2 (Thermo Fisher 14-9811-82, 1:250), PAX6 (Millipore AB2237, 1:250), TBR2 (Abcam ab183991, 1:1000), TUJ1 (Biolegend 801202, 1:1000), TBR1 (CST 49661S, 1:1000), CTIP2 (Absolute Antibody Ab00616-7.4, 1:500), RORβ (R&D Systems N7927, 1:100), LHX2 (Millipore ABE1402, 1:500), p53 (Leica CM5, 1:250), CC3 (CST 9661S, 1:250). Slides were mounted using either Vectashield (Vector Labs) or Fluoromount G (Thermo Fisher).

### Image acquisition and quantification

Images were captured using a Zeiss Axio Observer Z1 equipped with an apotome for optimal sectioning at 5x and/or 20x. For each experiment, 2-3 sections were imaged per brain. Images were captured with identical exposures. For image processing, maximum intensity projections were generated from Z-stacks and brightness was adjusted equivalently for all images using FIJI software. For quantification, images were cropped to 200 µm wide radial columns and cells were counted manually using FIJI cell counter, or automatically using QuPath (Bankhead et al. 2017). For QuPath quantification, parameters were adjusted as follows: requested pixel size = 0.1 µm, background radius = 5 µm, minimum area = 10 µm^2^, cell expansion = 2 µm, include cell nucleus and smoother boundaries boxes were unchecked. The threshold was set independently for each channel and maintained for each image in an experiment. For binning analysis, spatial coordinates for each detected cell were exported from either FIJI or QuPath and cells were assigned to one of five equally sized bins spanning from the ventricle to the pia to calculate the distribution.

### Live imaging

E12.5 cortices were dissected and dissociated as described above for SLAM-seq sample preparation with the following modifications: only one embryo was used per biological replicate, trypsinization was performed for 5 min, after resuspension in neural progenitor media, 175,000 cells were plated onto a poly-D-lysine treated 24-well glass-bottomed plate (MatTek). Cells were allowed to adhere to the plate for 2 hr, then images were captured every 10 min for 20 hr using a Zeiss Axio Observer Z1 microscope fitted with a Pecon incubation chamber as previously described (Mitchell-Dick et al. 2019). After 20 hr, cells were fixed in PFA for 20 min, permeabilized with 0.1% Triton-X100 for 10 min, blocked in 5% NGS for 30 min, incubated with primary antibody for 1 hr, and secondary antibody for 30 min. All immunostaining steps were performed at RT. The following primary antibodies were used: SOX2 (Thermo Fisher 14-9811-82, 1:1000), TBR2 (Abcam ab183991, 1:500), and TUJ1 (Biolegend 801202, 1:2000).

### Semi-cumulative labeling

Pregnant dams were intraperitoneally injected with BrdU (50 mg/kg) at t = 0 hr, followed by EdU (10 mg/kg) at t = 1.5 hr. Embryos were dissected at t = 2 hr. EdU was visualized using the Click-iT Plus Imaging Kit (Thermo Fisher C10638), followed by immunostaining using Ki67 (CST 12202, 1:250) and BrdU (Abcam ab6326, 1:200) primary antibodies as described above. Total cell cycle (Ts) and S-phase (Ts) were calculated as follows: S cells = EdU+; L cells = BrdU+EdU-; P cells = Ki67+; Ts = (S cells / L cells) * 1.5 hr; Tc = (P cells / S cells) * Ts

### Bulk RNA sequencing

E12.5 embryos were dissected in cold PBS and cortical tissue from single embryos were microdissected and flash frozen on dry ice. Three biological replicates were used per genotype. Tissue was homogenized in RLT buffer (Qiagen) and RNA was extracted using the RNeasy Plus Micro Kit (Qiagen). Libraries were prepared using the TruSeq Stranded mRNA kit (Illumina) and sequenced on the NovaSeq platform with 150 bp paired-end reads. Read quality was assessed with FastQC and adapter trimming was performed using Cutadapt. Reads were aligned to the mm39 genome using STAR (Dobin et al. 2013). Mapped reads were counted and assigned to genes using featureCounts (Liao, Smyth and Shi 2014), and differential expression in cKO and dcKO versus control samples was determined using DESeq2 (Love, Huber and Anders 2014).

### Transcriptional shut-off

E12.5 cortices were dissected and dissociated as described above for SLAM-seq sample preparation with the following modifications: only one embryo was used per biological replicate, trypsinization was performed for 5 min, after resuspension in neural progenitor media each replicate was split into three wells of a poly-D-lysine coated 24-well plate. At t = 0, actinomycin D (Sigma A9415) was added to the media to a final concentration of 5 µg/mL. Cells were lysed at t = 0, 2, 8 hr by addition of Trizol (Thermo Fisher 15596026) and RNA was extracted according to manufacturer’s instructions. cDNA was generated from 500 ng total RNA using iScript cDNA Synthesis Kit (Bio-Rad). qPCR was performed using iTaq SYBR Green Supermix (Bio-Rad). qPCR primers are listed in Supplemental Table S6. Decay data was fitted to the single exponential decay equation (y = y_o_ * e^-kt^) using the nls function in R to determine the degradation rate constant, k. Half-life was calculated as t_1/2_ = (ln(2))/k.

### Statistical analysis

Sample collection, data acquisition, and quantifications were performed blind with respect to genotype. Statistical tests, *p*-values and sample size for each analysis are reported in Supplemental Table S7.

### Data availability

Sequencing data will be deposited on GEO prior to publication.

## Acknowledgements

We thank members of the Silver and Hu labs for helpful discussions. This work was supported by the Extramural and Intramural Research Programs of the National Institutes of Health: F32HD107972 to L.D.S., R01NS083897, R01NS120667, R01NS110388, R01MH132089, R21NS128374 to D.L.S., and Z01ES102745 to G.H.

## Author Contributions

L.D.S. and D.L.S. conceived the study. L.D.S., D.L.S., and G.H. designed experiments. L.D.S., J.R.E., B.L., B.D.B. performed experiments, and analyzed data. L.D.S. and D.L.S wrote the manuscript. All authors edited the manuscript.

---

**Supplemental Figure 1.**
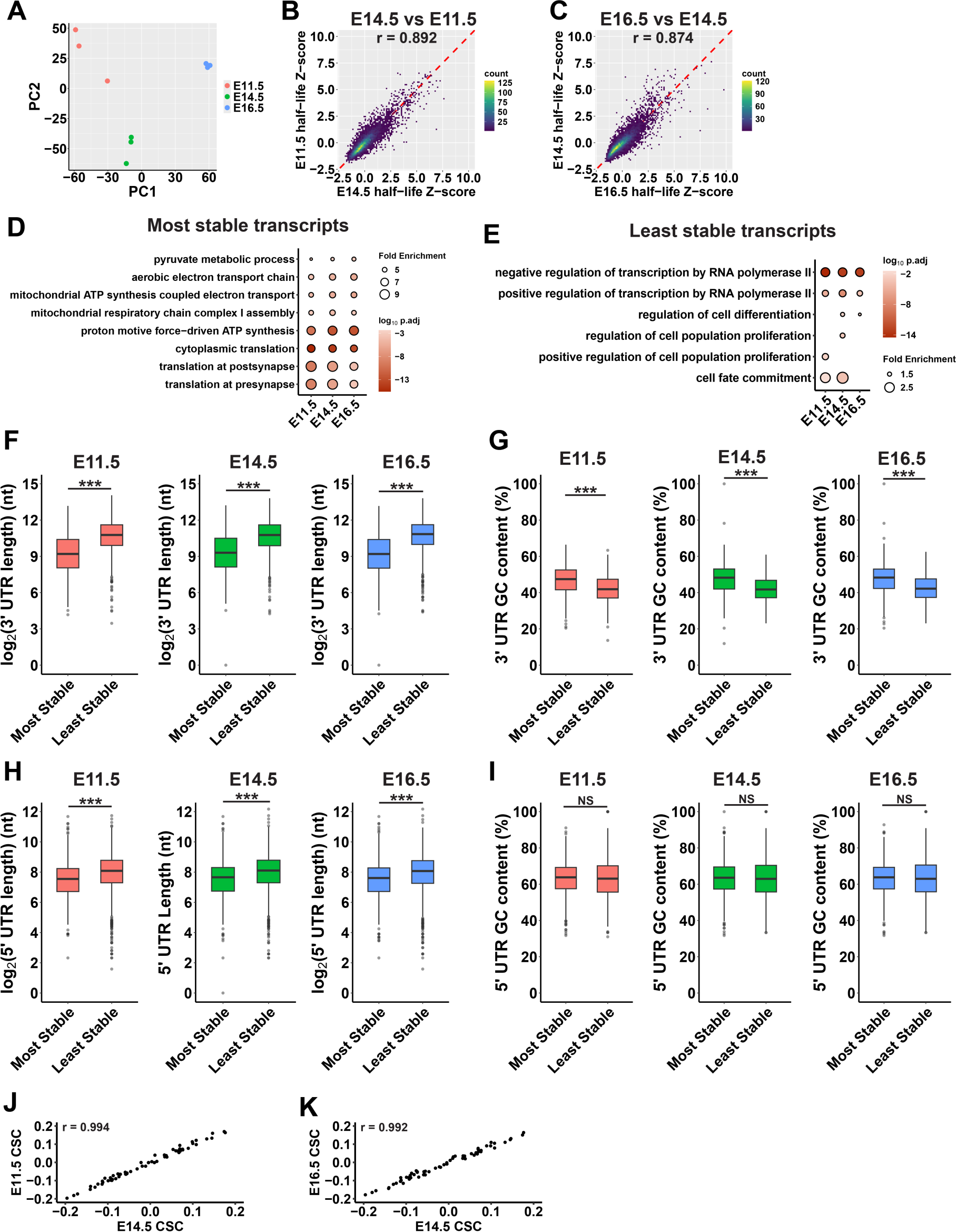
Features of mRNA stability in the developing cortex. (A) Principal component analysis of biological replicates from SLAM-seq half-life data. (B) Correlation between z-score normalized half-lives at E11.5 and E14.5. Red dashed line indicates y = x line. r represents the Pearson correlation coefficient. (C) As in B, for E14.5 and E16.5. (D and E) GO analysis of top 10% most stable and least stable transcripts. Circle size represents fold enrichment and color represents adjusted *p* value. (F) Comparison of 3’ UTR lengths of the top 10% most and least stable transcripts at each stage. (G) As in E, comparing percentage GC content. (H, I) As in F, G, for 5’ UTRs. (J) Comparison of CSC values between E11.5 and E14.5. r represents the Pearson correlation coefficient. (K) As in J, for E14.5 and E16.5. \*\*\**p* < 0.001. Wilcoxon rank-sum test (F, H), Welch’s two-sample t-test (G, I).

**Supplemental Figure 2.**
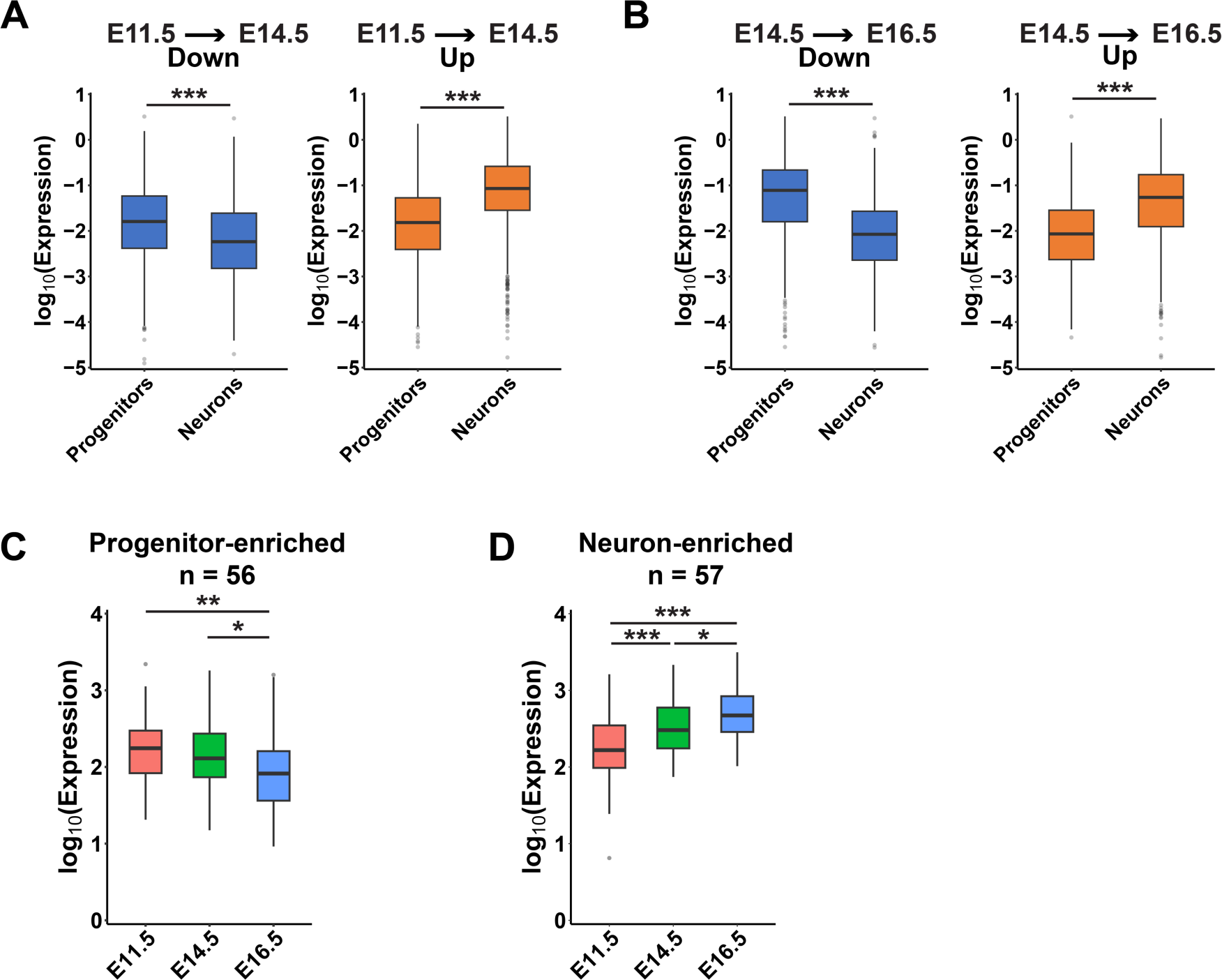
Developmentally regulated genes have cell type enriched expression patterns. (A) Expression of differentially expressed genes at E14.5 compared to E11.5. Expression data from publicly available in scRNA-seq (Di Bella et al. 2021). (B) As in A, for differentially expressed genes at E16.5 compared to E14.5. (C) Expression of progenitor-enriched genes across development using SLAM-seq expression data from this study. (D) As in C, for neuron-enriched genes. \*\*\**p* < 0.001. Wilcoxon rank-sum test (A, B), one-way ANOVA with Tukey’s HSD post-hoc test (C, D).

**Supplemental Figure 3.**
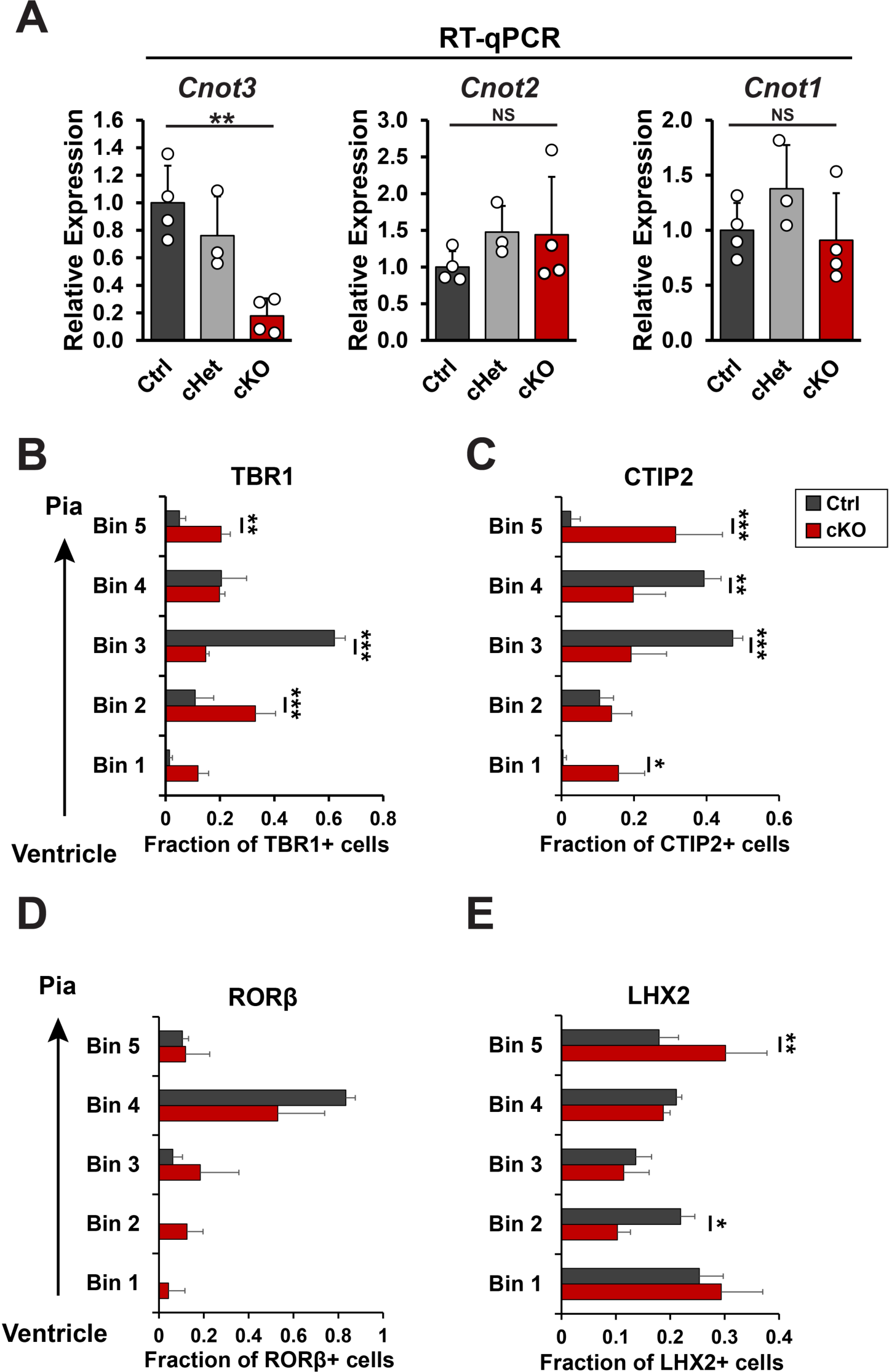
C*n*ot3 is required for neuronal lamination. (A) Expression of *Cnot1, Cnot2, and Cnot3* in E12.5 cortical lysates measured by RT-qPCR. (B-D) Distribution of the indicated marker at E18.5 across five equally sized cortical bins spanning the ventricular surface (Bin 1) to the pia (Bin 5). \**p* < 0.05, \*\**p* < 0.01, \*\*\**p* < 0.001. One-way ANOVA with Tukey’s HSD post-hoc test (A), two-way ANOVA with Tukey’s HSD post-hoc test (B-D). Error bars represent standard deviation.

**Supplemental Figure 4.**
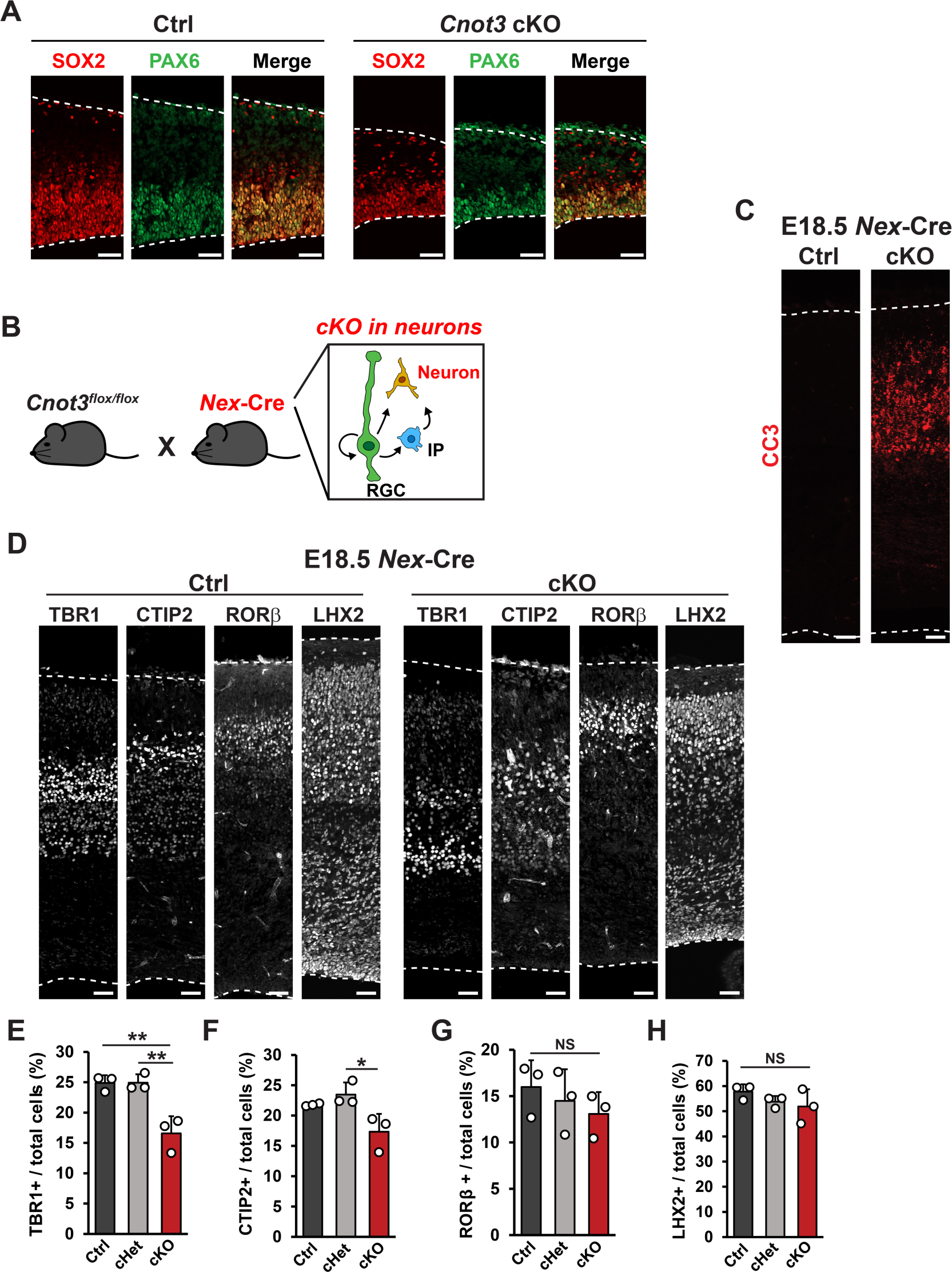
C*n*ot3 alters RGC distribution and is required for survival of post-mitotic neurons. (A) Immunofluorescence against SOX2 and PAX6 in E14.5 control and *Emx1*-Cre cKO cortices. (B) Schematic showing strategy for *Cnot3* cKO in neurons using *Nex*-Cre. (C) Immunofluorescence against CC3 in E18.5 control and *Nex*-Cre cKO cortices. (D) Immunofluorescence against the indicated marker in E18.5 control and *Nex*-Cre cKO cortices. (E-H) Quantification of density for the indicated markers (n = 3 embryos per genotype). \*\**p* < 0.01. One-way ANOVA with Tukey’s HSD post-hoc test (F-H). Error bars represent standard deviation. Scale bars: 50 µm for all images.

**Supplemental Figure 5.**
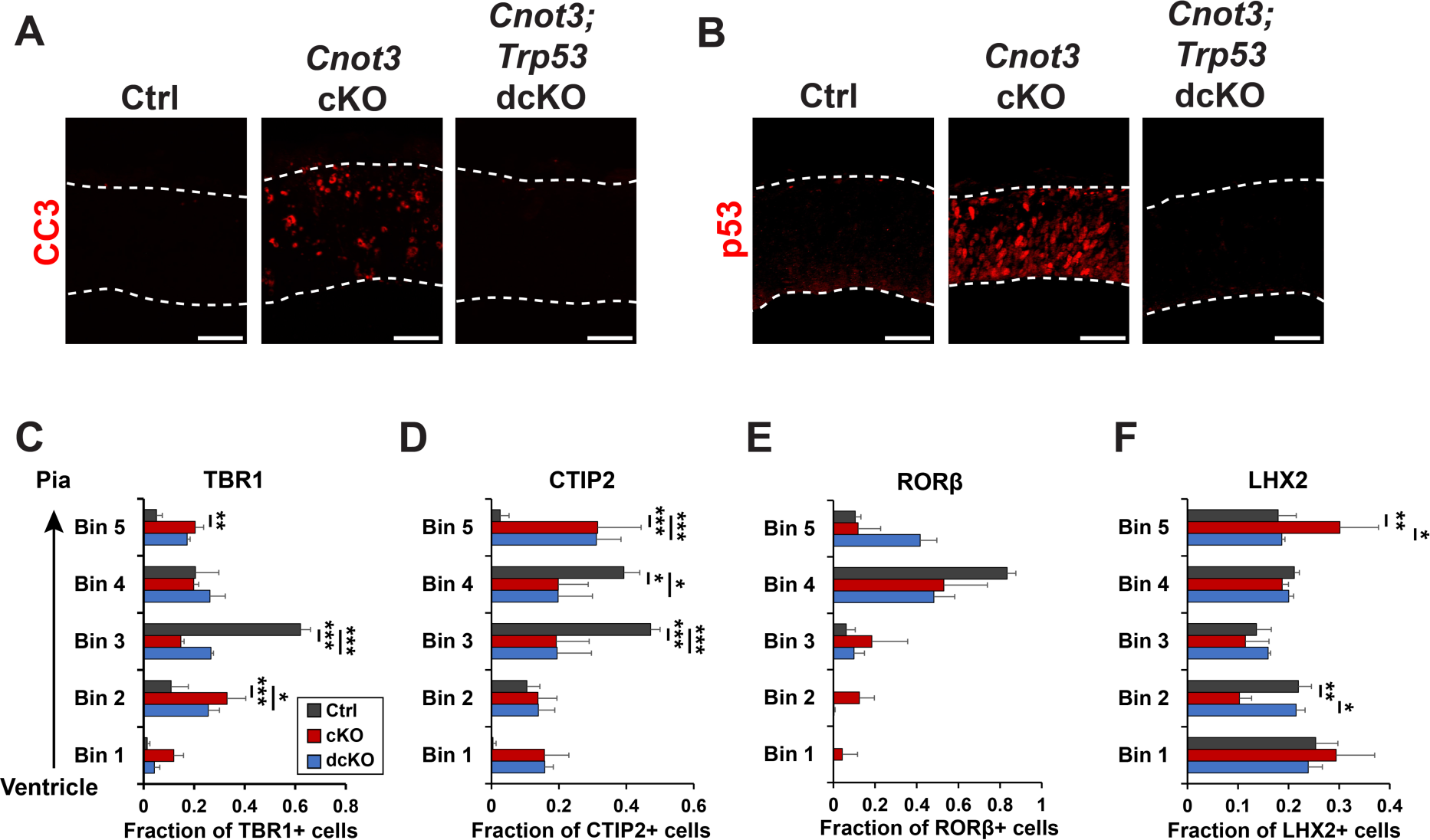
C*n*ot3*;Trp53* dcKO brains have no apoptosis but exhibit some altered lamination patterns. (A) Representative images of Immunofluorescence against CC3 in E12.5 cortices showing rescue of apoptosis in dcKO mice. (B) Representative images of immunofluorescence against p53 in E12.5 cortices showing rescue of p53 accumulation in dcKO mice. (C-F) Distribution of the indicated marker at E18.5 across five equally sized cortical bins spanning the ventricular surface (Bin 1) to the pia (Bin 5). \**p* < 0.05, \*\**p* < 0.01, \*\*\**p* < 0.001. Three-way ANOVA with Tukey’s HSD post-hoc test. Error bars represent standard deviation. Scale bars: 50 µm (A, B).

**Supplemental Figure 6.**
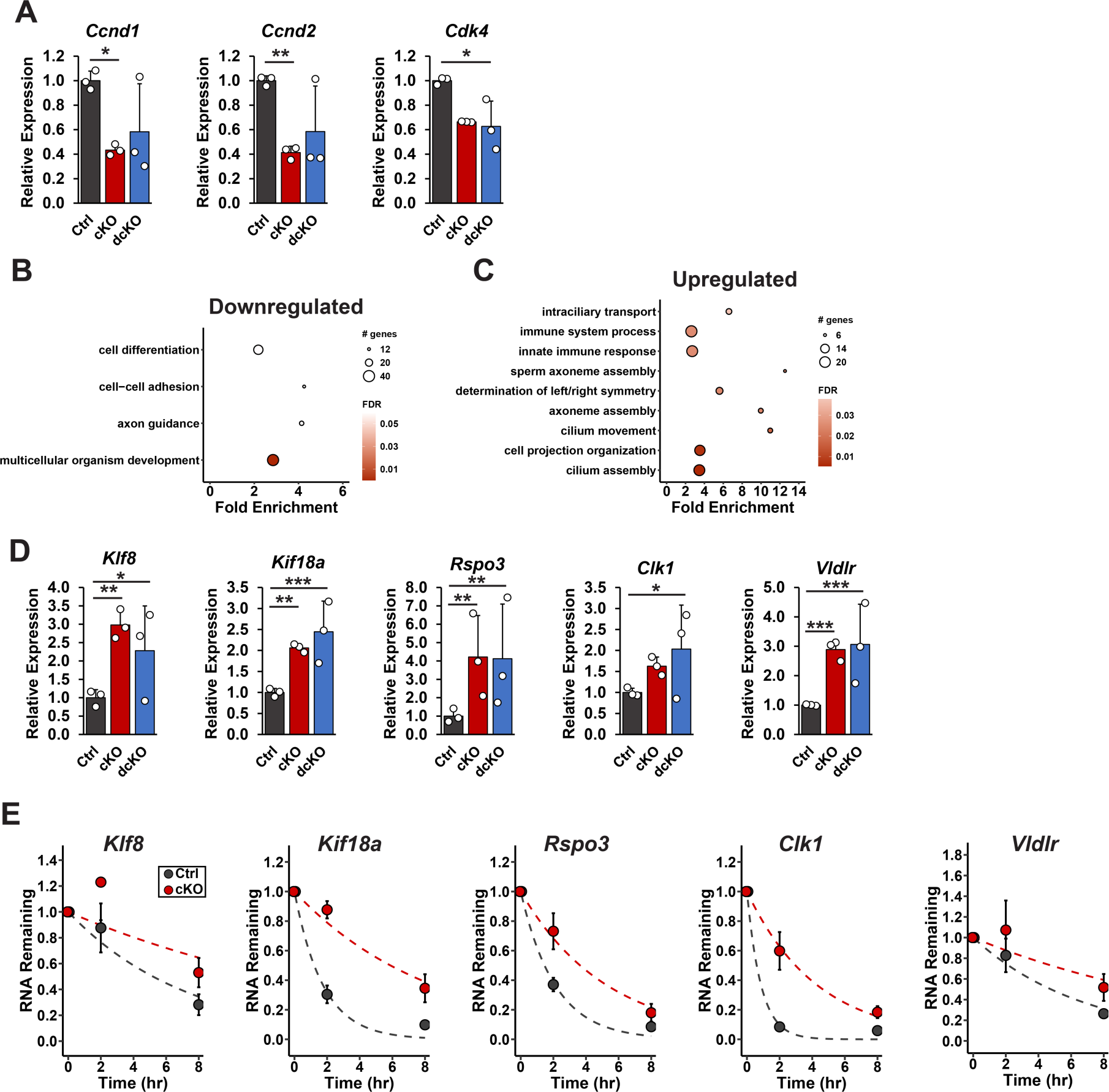
C*n*ot3-dependent regulation of gene expression. (A) Expression of the indicated cell cycle gene at E12.5 measured by RNA-seq. (B, C) GO analysis of transcripts that are downregulated (B) or upregulated (C) in both cKO and dcKO cortices at E12.5. Circle size corresponds to number of genes and color represents FDR. (D) Transcript levels at E12.5 measured by RNA-seq. (E) Decay curves measured by RT-qPCR following transcriptional shut-off in E12.5 primary cultures. Dashed lines indicate fit to exponential decay equation. \**p* < 0.05, \*\**p* < 0.01, \*\*\**p* < 0.001. Values shown are adjusted *p*-values from DESeq2 (A, D). Error bars represent standard deviation.

## List of Supplemental Tables

Supplemental Table 1: SLAM-seq half-life and expression data for E11.5, E14.5 and E16.5

Supplemental Table 2: List of transcripts with variable half-lives: related to Fig. 2

Supplemental Table 3: Differential expression analysis between developmental stages

Supplemental Table 4: List of genes enriched in progenitors or neurons

Supplemental Table 5: Differentially expressed genes in E12.5 cKO and dcKO cortices

Supplemental Table 6: Primers used in this study

Supplemental Table 7: List of statistical analyses

